# General anaesthesia disrupts complex cortical dynamics in response to intracranial electrical stimulation in rats

**DOI:** 10.1101/2020.02.25.964056

**Authors:** A. Arena, R. Comolatti, S. Thon, A.G. Casali, J.F. Storm

## Abstract

The capacity of the human brain to sustain complex dynamics consistently drops when consciousness fades. Several recent studies in humans found a remarkable reduction of the complexity of cortical responses to local stimulation during dreamless sleep, general anaesthesia, and coma. So far, this perturbational complexity has never been estimated in non-human animals *in vivo*. Here, we quantify the complexity of electroencephalographic responses to intracranial electrical stimulation in rats, comparing wakefulness to propofol, sevoflurane, and ketamine anaesthesia. We confirm the changes previously observed in humans: from highly complex evoked activity during wakefulness, to simpler responses, suppression of high frequencies, and reduced phase-locking with propofol and sevoflurane. We then deepen our mechanistic understanding by analyzing functional connectivity, and by showing how these parameters dissociate with ketamine, and depend on intensity and site of stimulation. This approach opens the way for further direct investigations of the mechanisms underlying brain complexity and consciousness.

## INTRODUCTION

A longstanding challenge in neuroscience has been the identification of robust and neuronally based measures of consciousness. Recently, brain complexity, defined as the combination of functional differentiation and integration in thalamocortical systems, has gained growing attention as promising candidate^1,2,3^. For example, a reliable association between conscious states and complexity of global network dynamics has been demonstrated by means of both electroencephalography (EEG)^4,5^ and imaging of spontaneous brain activity^6,7^. A highly accurate method to assess complexity of causal, cortical dynamics is the Perturbational Complexity Index (PCI), which was originally introduced and validated for discrimination between unconscious and conscious unresponsive patients^1,8^. PCI is based on a perturbational approach: brief transcranial magnetic stimulation (TMS) or intracranial electrical stimulation are used to trigger cortical activity, and the spatiotemporal complexity of the EEG-recorded event related potentials (ERPs) is quantified. Long-lasting responses that are both temporally differentiated and distributed among cortical areas (high PCI) are evoked whenever subjects have conscious experiences, such as during wakefulness, rapid eye movement (REM) sleep or ketamine anaesthesia, when dreams or hallucinations occur^1,9,8,10,11^. In contrast, during dreamless non-REM (NREM) sleep, general anaesthesia, and unresponsive wakefulness syndrome (UWS, or “vegetative state”), the ERPs appear to be either local and short-lasting (less integrated), or global and stereotyped (less differentiated), yielding low PCI^1,9,8,10,11^.

Although the PCI method has been thoroughly tested in humans, it has so far not been tested in any non-human species *in vivo* and it is not clear to what extent rodent brains can sustain complex dynamics in response to brief direct stimulation. Developing animal models and extending the application of PCI is of paramount importance, since the neuronal mechanisms underlying the engagement or interruption of complex cortical activations are still uncertain, and mechanistic investigations may lead to intervention strategies for restoring complexity and awareness in brain-injured, unconscious patients. Here we implement and test the PCI method in rats. By measuring the complexity of intracranial EEG responses to cortical stimulation, we show that the awake rat brain supports complex cortical activations, which turn into simplified, less integrated responses during general anaesthesia. We further demonstrate that the disruption of long-lasting complex activations are associated with suppression of high-frequency EEG power and reduced phase-locking, supporting the hypothesis that neuronal hyperpolarization might prevent cortical neurons from engaging in durable, complex interactions^12,13,14,15^.

## RESULTS

### Single pulse electrical stimulation triggered complex ERPs during wakefulness, but not during propofol anaesthesia

We recorded epidural EEG activity from 16 screw electrodes chronically implanted through the skull in head- and body-restrained male, adult rats. Recording electrodes were in contact with the dura and organized in a symmetric grid, covering most of the cortex in both hemispheres (M2, secondary motor cortex; M1, primary motor cortex; S1, primary somatosensory cortex; RS, retrosplenial cortex; PA, parietal associative cortex; V1, primary visual cortex; V2, secondary visual cortex; GND, ground electrode over cerebellum). We stimulated right M2 by single monophasic, electrical current pulses (typically: 1 ms duration, 50 µA amplitude, at 0.1 Hz) via a chronically implanted bipolar electrode, located 4.38 ± 0.26 mm rostral from bregma, 0.47 ± 0.09 mm below the cortical surface, mainly corresponding to layer II/III (based on histology after recording, 8 rats; **Fig. 1a**; all values are reported as mean ± SEM). Pulse trains delivered at similar coordinates triggered coordinated whisker deflections^16^, whereas EEG responses following single stimuli were not measurably contaminated by movements (**Supplementary Fig. 1, Video 1-2**) and were reproducible throughout recording sessions and across days (**Supplementary Fig. 2**). No correlation between the stimulating electrode locations and ERP amplitude or duration was found (**Supplementary Fig. 3**).

**Fig. 1.**
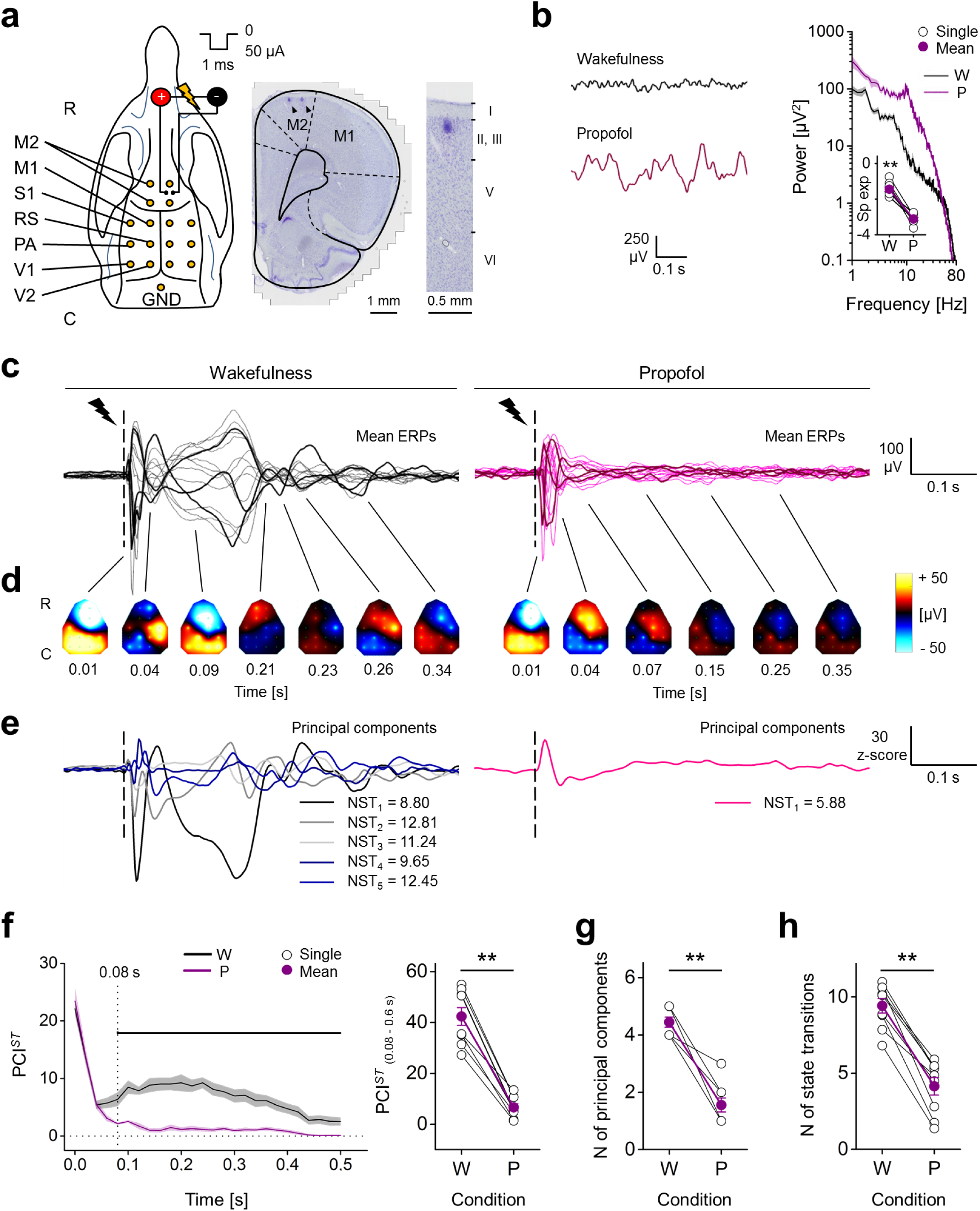
Spatiotemporal dynamics of evoked responses to electrical stimulation of M2 during wakefulness and propofol anaesthesia. **a**, *Left*, Positions in the rat skull of the16 screw electrodes (yellow dots) and bipolar stimulating electrode used for recording EEG and triggering ERPs (R-C: rostral-caudal). *Right*, coronal brain section (Nissl staining) showing the location of stimulating electrode in right M2. Black arrowheads indicate marks from the 2 poles of the bipolar electrode. *Far right*, magnified view showing the site of one pole relative to cortical layers. **b**, *Left*, spontaneous EEG from one rat during wakefulness (W) and propofol (P) anaesthesia. *Right*, mean periodograms of spontaneous activity from one animal in same conditions (shades represent SEM), and spectral exponents from all rats (*inset*). **c-e**, EEG responses to single pulse electrical stimulation (1 ms, 50 µA; dashed line) from one rat during wakefulness and propofol anaesthesia. **c**, Butterfly plots show superimposed mean ERPs from all recording electrodes (ERPs from 3 channels are in bold for clarity) and **d**, their spatial distributions at different time points (interpolated ERPs, color-coded). **e**, Derived principal components (from the same data as in **c, d**) with corresponding numbers of state transitions (NST). **f**, *Left*, time course of PCI^*ST*^ averaged from 9 rats in wakefulness and propofol anaesthesia (0 s: stimulus onset; shades represent SEM; horizontal line indicates statistical difference, *P* < 0.05). *Right*, PCI^*ST*^ quantified within the time window 0.08-0.60 s. **g**, Number of principal components and **h**, average numbers of state transitions across conditions for all rats.

We performed electrophysiological recordings in 9 rats during wakefulness and propofol anaesthesia (∼1.1 mg/kg/min, i.v.) at a depth that produced spontaneous, slow, high-amplitude EEG oscillations and was sufficient to abolish any detectable motor response to pain stimuli. The redistribution of EEG power from high to low frequencies was confirmed by a reduced spectral exponent of the periodrogram (range: 20-40 Hz) from wakefulness to propofol anaesthesia (**Fig. 1b**; wakefulness: −1.44 ± 0.12, propofol: −3.12 ± 0.09; Wilcoxon S-R test, *P* = 0.004). During wakefulness, single pulse stimulation (1 ms, 50 µA) triggered long-lasting ERPs, including an early, fast, high-voltage response followed by multiple changes in polarity over time and across cortical areas. During anaesthesia, however, the same stimulation produced only a similar initial activation, followed by fewer polarity changes (**Fig. 1c-d, Supplementary Video 3-4**). The ERP complexity was quantified by Perturbational Complexity Index state-transition (PCI^*ST*^), a version of PCI based on the state transitions of principal components of the EEG response^10^ (**Fig.1e**; see Methods). We initially assessed the PCI^*ST*^ time course, using sliding windows of 0.1 s. Immediately after stimulation, PCI^*ST*^ was similar across conditions and quickly decayed. Soon afterwards, however, complexity (PCI^*ST*^) built up reaching a maximum at 0.21 ± 0.04 s during wakefulness, and differed significantly from propofol anaesthesia starting from 0.08 s until ERP ended (**Fig. 1f;** Wilcoxon S-R test). Thus, in the time window 0.08-0.6 s, PCI^*ST*^ showed a clear reduction from wakefulness to propofol anaesthesia, for both single rats and the population (**Fig. 1f;** wakefulness: 42.35 ± 3.47, propofol: 6.63 ± 1.38; Wilcoxon S-R test, *P* = 0.004). In the same time range, the reduced PCI^*ST*^ was determined both by a reduced number of principal components (**Fig. 1g;** wakefulness: 4.44 ± 0.18, propofol: 1.55 ± 0.24; Wilcoxon S-R test, *P* = 0.004) and a reduced number of state transitions over time (**Fig. 1h;** wakefulness: 9.42 ± 0.48, propofol: 4.14 ± 0.57; Wilcoxon S-R test, *P* = 0.004; **Supplementary Fig. 4**).

### An OFF period preceded the early interruption of deterministic response during propofol anaesthesia

Next, we quantified the changes in spectral power caused by the stimulation, within a high frequency range (HF, 20-40 Hz) that has been shown to maximize the difference between active and silent (OFF) periods of a neuronal network that oscillates between depolarized and hyperpolarized states^17,18,19,20^, as typically occurs during NREM sleep and general anaesthesia^19,20,21^. We also considered the responses as “deterministic”, i.e. reliably driven by the stimulation, if the evoked potentials were largely reproducible in phase when the same stimulation was repeated^22^. Consequently, the duration of the deterministic neuronal response was defined as the last time point of the phase-locked component of the ERP measured as “inter-trial phase clustering” (ITPC)^23^, in the 8-40 Hz frequency range^12,24,10^.

Coherently with the PCI^*ST*^ time course, the ERP showed an early, transient increase in HF power during both wakefulness and propofol anaesthesia. A later HF activation, sustained until the end of the response in all channels, was detected only during wakefulness. By contrast, propofol anaesthesia induced a deep HF suppression in most of the channels at 0.08 ± 0.01 s (**Fig. 2a-b**). When averaging the relative HF power during the suppression (0.08-0.18 s), across channels and rats, we found that it decreased from wakefulness to propofol anaesthesia, to values below the baseline in all rats, indicating an OFF period (**Fig. 2c**; wakefulness: 4.13 ± 0.59 dB, propofol: −0.79 ± 0.25 dB; Wilcoxon S-R test, *P* = 0.004). Traces of HF suppression after stimulation were observed also during wakefulness, but these were briefer and shallower than with propofol (**Supplementary Fig. 5**), and were not seen after averaging (**Fig 2b-c**). Moreover, during wakefulness the stimulation evoked durable, phase-locked responses in all electrodes. In contrast, during propofol anaesthesia the OFF period preceded an earlier drop of ITPC in all channels, followed by transient and not phase-locked HF activations in few cortical areas (**Fig. 2a-b**). Consequently, the phase-locked response, measured by averaging the ITPC drop time across channels and rats, was significantly briefer during propofol (0.13 ± 0.01 s) than during wakefulness (0.32 ± 0.04 s; Wilcoxon S-R test, *P* = 0.004; **Fig. 2d**).

**Fig. 2.**
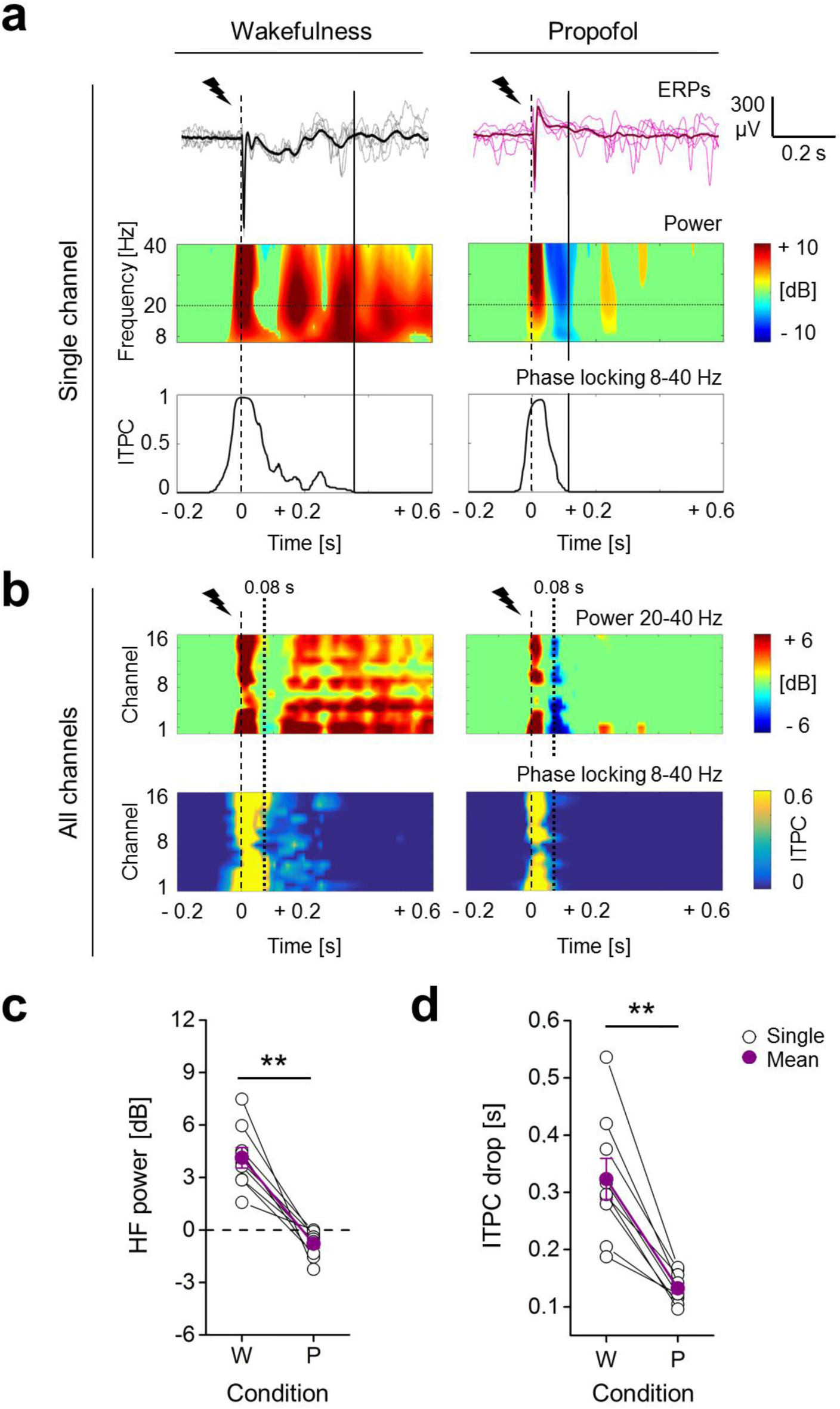
Propofol anaesthesia induced suppression of high frequencies and reduced phase-locking in response to electrical stimulation, compared to wakefulness. **a**, Example of epidural EEG response to single pulse electrical stimulation (1 ms, 50 µA; dashed line) from the same rat during wakefulness (*left*) and propofol anaesthesia (*right*). The mean ERPs (bold) and 5 consecutive single trials from the same frontal channel (M2) are shown in both conditions (*top)* with relative spectrogram (*middle)* and ITPC (*below*) averaged in the frequency range 8-40 Hz. The continuos vertical lines indicate the time point of the ITPC drop. **b**, Time course of the average HF power in the 20-40 Hz range (*top*), and the averaged ITPC in the 8-40 Hz range (*below*) plotted for all channels from the same rat and conditions of **a**. The dotted vertical line at 0.08 s indicates the mean onset of HF suppression across rats during propofol anaesthesia. **c**, The mean HF power (in time range: 0.08-0.18 s) and (**d)** the duration of phase-locking across trials (time of ITPC drop) for all animals (n = 9) during wakefulness (W) and propofol anaesthesia (P).

Next, we increased the stimulation intensity from 40 to 100 µA (1 ms; 5 rats), attempting to compensate for the inhibiting effect of propofol^25,26^ (**Fig. 3a, Supplementary Fig. 6**). The resulting neuronal excitation, quantified by the root mean squared (rms) amplitude of the first deflections of mean ERPs (Early ERP rms, up to 0.05 s), increased linearly with stimulus intensity, during both wakefulness (**Fig. 3b**; Friedman test, *P* = 0.002; linear fit, *P* = 0.006, R^2^ = 0.988) and propofol anaesthesia (Friedman test, *P* = 0.002; linear fit, *P* = 0.016, R^2^ = 0.968). Overall, no significant differences in Early ERP rms were detected between wakefulness and anaesthesia (Friedman test, *P* = 0.917). With propofol, the increased excitation was accompanied by a deeper HF suppression as indicated by the linear decrease of HF power as function of stimulus intensity (**Fig. 3c**; Friedman test, *P* = 0.005; linear fit, *P* = 0.015, R^2^ = 0.969). In contrast, during wakefulness, the HF power was always above baseline, thus higher than during propofol (Friedman test, *P* = 1.767*10^−7^), and no change with stimulus intensity was detected (Friedman test, *P* = 0.178). During propofol anaesthesia, higher stimulus intensities were also linearly related to prolonged OFF periods (**Fig. 3d, Supplementary Fig. 5**; Friedman test, *P* = 0.005; linear fit, *P* = 0.024, R^2^ = 0.953). In contrast, we did not found intensity dependent change in the brief HF suppressions during wakefulness (**Supplementary Fig. 5**). When using these changes in duration to assess a temporal relation between the OFF period and the interruption of ITPC with propofol (averaging across rats for each channel and stimulation intensity), we found a highly significant correlation between the time points of the OFF period end and the ITPC drop (**Fig 3d**; linear fit, *P* = 6.079*10^−8^, R^2^ = 0.384). PCI^*ST*^ was always higher and ITPC more long-lasting during wakefulness than with propofol, regardless of the stimulus intensity, in all tested animals (**Supplementary Fig. 6**).

**Fig. 3.**
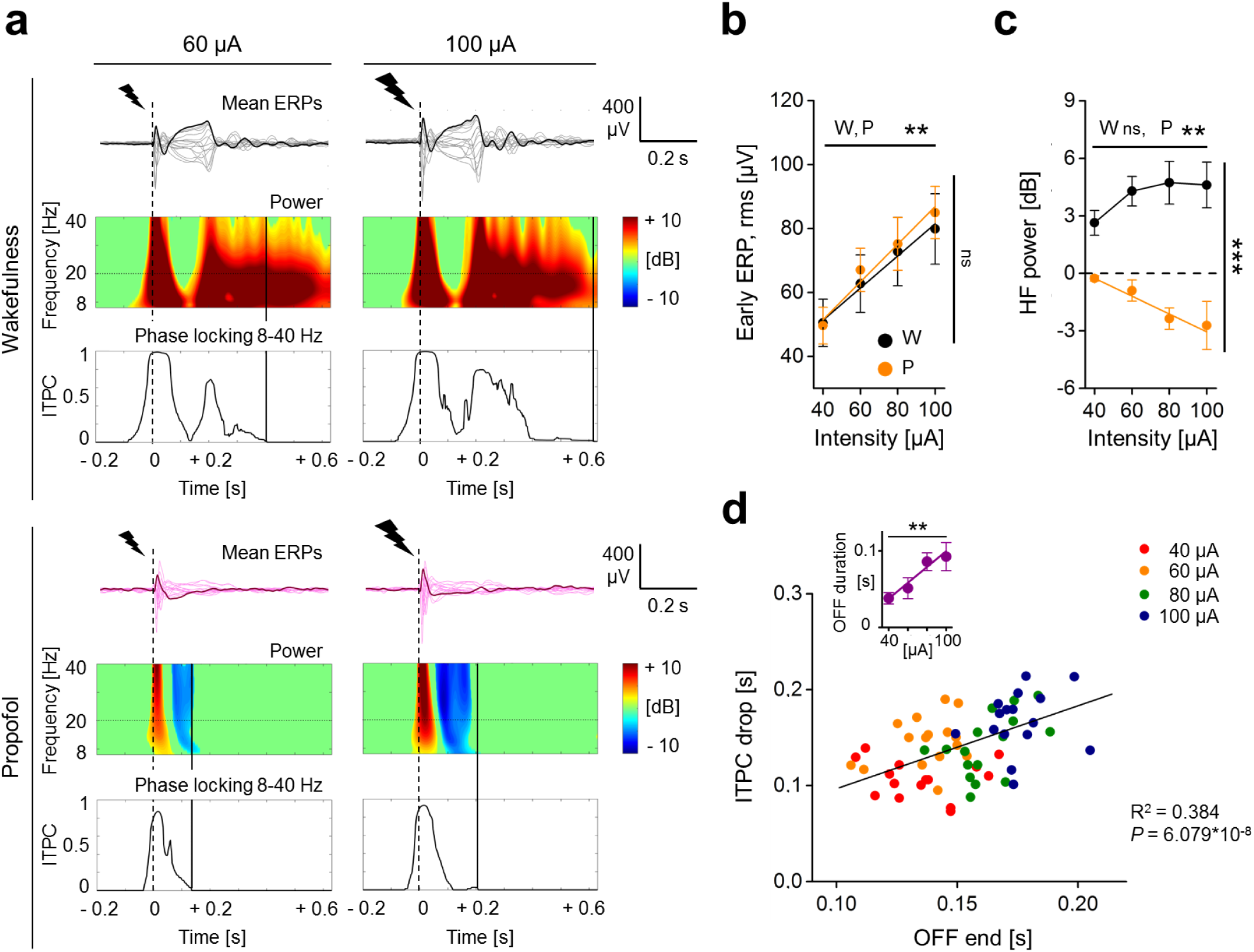
During propofol anaesthesia, the HF suppression was deeper and lasted longer after stronger stimulation, and correlated in time with the drop in phase-locking. **a**, Mean ERPs from the same rat in response to single pulse electrical stimulations (dashed lines) of two different intensities (60 µA *left*, 100 µA *right*), during wakefulness (*top*) and during propofol anaesthesia (*bottom*). The butterfly plots show averaged ERPs from all recording electrodes superimposed, with one mean ERP from the same occipital channel (V1) shown in bold for clarity. *Below*, the spectrogram and the ITPC plots (8-40 Hz) for the highlighted channel are shown. The continuous vertical lines indicate the time points of ITPC drop. **b**, Quantification of early ERP rms amplitude (up to 0.05 s) and **c**, HF power (in range 0.08-0.18 s) as a function of increasing stimulation intensity during both wakefulness (W) and propofol anaesthesia (P). Values are averaged across channels and animals (n = 5). **d**, During propofol anaesthesia, the time spent with suppression of HF power (OFF duration) increased as a function of stimulus intensity (*inset*), and there was a positive correlation between the latency of the end of the OFF period (time of OFF end) and the duration of phase-locking across trials (time of ITPC drop). The averaged values across rats for each channel and stimulus intensity (color-coded) are plotted. The coefficient of determination R^2^ and the *P* value are reported.

### ERPs during ketamine anaesthesia showed intermediate complexity

In order to test whether the PCI^*ST*^ reduction with propofol might be related to behavioural unresponsiveness per se, we repeated single pulse stimulations (1 ms, 50 µA; 8 rats) during ketamine anaesthesia, which was found to maintain high brain complexity in humans^9^, at a dose that abolished all motor responses to painful stimuli (∼1.8 mg/kg/min i.v; **Fig. 4a, Supplementary Video 5**). Similarly to wakefulness, the spontaneous EEG activity with ketamine showed fast, shallow oscillations, with similar spectral exponent (**Fig 4b;** 7 rats; wakefulness: −1.21 ± 0.24, ketamine: −0.97 ± 0.09; Wilcoxon S-R test, *P* = 0.156). The PCI^*ST*^ time course revealed a similar initial complexity of the ERP across conditions that quickly decayed. Like wakefulness, but unlike propofol, during ketamine anaesthesia PCI^*ST*^ increased soon after the initial decay, reaching a peak at ∼0.2 s and gradually decreased until the ERP end. From 0.08 s the complexity level with ketamine was lower than during wakefulness, but transient periods of similar PCI^*ST*^ were detected (**Fig. 4c;** Wilcoxon S-R test). Similarly, from 0.16 s, PCI^*ST*^ with ketamine was significantly higher than during propofol anaesthesia and only brief periods of similar complexity were identified until the end of ERP (**Fig. 4c;** Mann-Whitney test). Coherently, PCI^*ST*^ differed between wakefulness and ketamine anaesthesia for the 0.08-0.6 s period (**Fig. 4c;** wakefulness: 41.80 ± 5.17, ketamine: 21.14 ± 4.48; Wilcoxon S-R test, *P* = 0.008), but also between ketamine and propofol anaesthesia (**Fig. 4c;** Mann-Whitney test, *P* = 0.024). The intermediate PCI^*ST*^ with ketamine was explained by a similar number of EEG components compared to wakefulness (**Fig. 4d;** wakefulness: 4.37 ± 0.18, ketamine: 3.62 ± 0.62; Wilcoxon S-R test, *P* = 0.25), with a reduced number of state transitions (**Fig. 4e;** wakefulness: 9.36 ± 0.81, ketamine: 5.50 ± 0.31; Wilcoxon S-R test, *P* = 0.008; **Supplementary Fig.4**). After the similarly complex initial response, also with ketamine an OFF period occurred at 0.08 ± 0.01 s (**Fig. 4f**; wakefulness: 4.25 ± 0.78 dB, ketamine: - 1.47 ± 0.44 dB, Wilcoxon S-R test, *P* = 0.008), but no consistent changes in ITPC drop time were found (**Fig. 4g**; wakefulness: 0.32 ± 0.04 s, ketamine: 0.23 ± 0.04 s, Wilcoxon S-R test, *P* = 0.195). A late increase in HF power after the OFF period occurred in a higher proportion of channels during ketamine anaesthesia than with propofol (**Fig. 4h**; propofol, 0.33 ± 0.07; ketamine, 0.90 ± 0.07; Mann-Whitney test, *P* = 0.001), while in wakefulness, it was seen in all channels in all 12 animals. We then assessed whether this resumption of HF activity was still associated with the deterministic neuronal activation by comparing the time of ITPC drop with the onset of late HF power increment. Only channels that presented a late increase in HF power were included for this comparison. During wakefulness, the resumption of HF activity was largely deterministic, occurring within a period of significant ITPC (**Fig. 4i**; Wilcoxon S-R test, *P* = 4.883*10^−4^), whereas during propofol the late HF power was not phase-locked as it occurred after the fading of ITPC (**Fig. 4i**; Wilcoxon S-R test, propofol, *P* = 0.008). Coherently with the intermediate complexity level during ketamine, the late HF activity observed in this condition occurred with a variable relationship with respect to the drop of ITPC (**Fig. 4i**; Wilcoxon S-R test, *P* = 1).

**Fig. 4.**
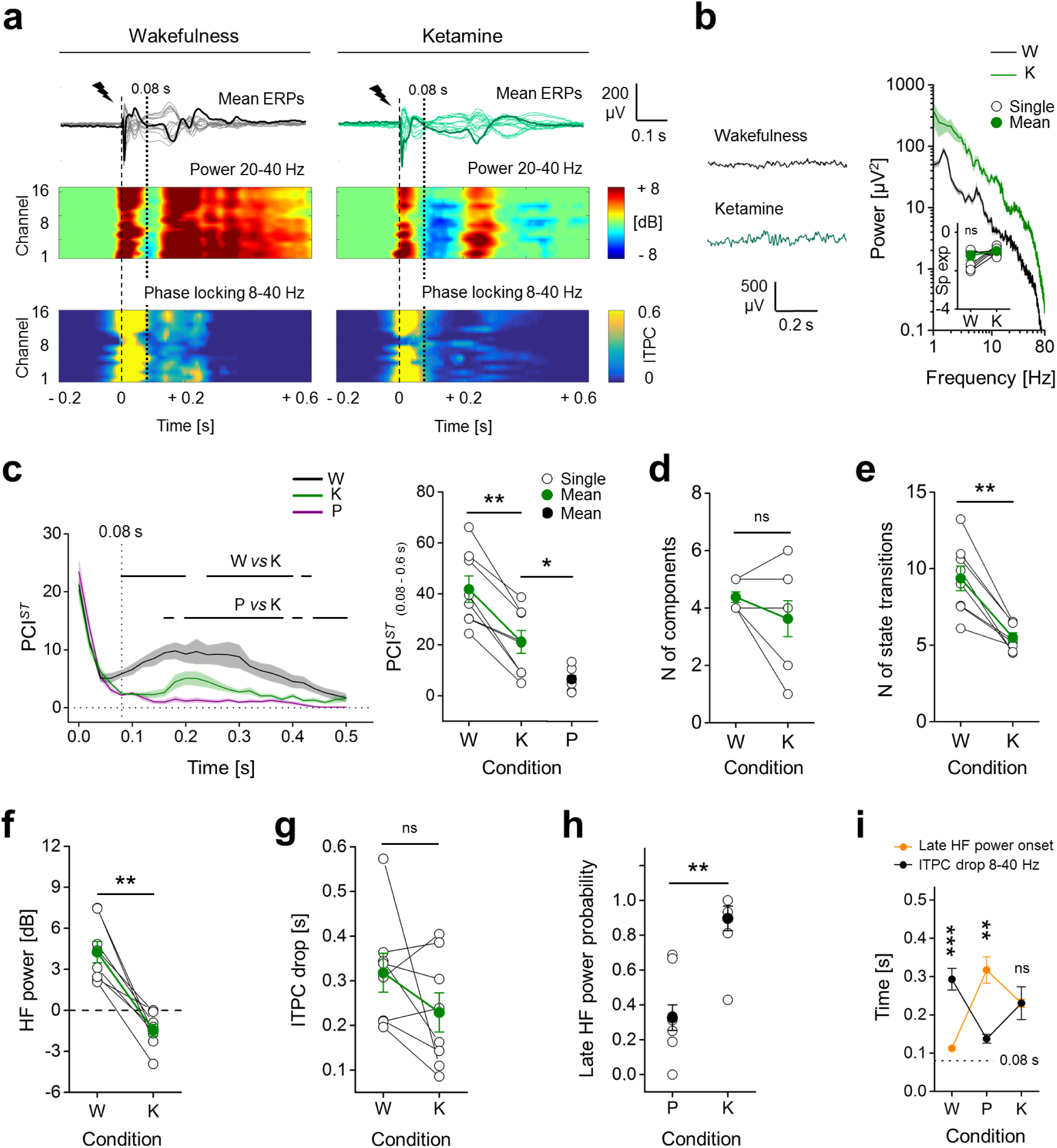
ERPs with ketamine showed intermediate PCI^*ST*^, with OFF period, but sustained ITPC. **a**, Mean ERPs from all electrodes in response to single pulse stimulation (1 ms, 50 µA; dashed line) shown superimposed, from the same rat during wakefulness and ketamine anaesthesia. One averaged ERP from the same channel (M2) is in bold for clarity. Spectrograms of HF power and ITPC for all channels are shown below. Vertical dotted line at 0.08 s indicates the average time of OFF period onset. **b**, Spontaneous EEG (*left*) and relative mean periodograms (shades represent SEM; *right*) are shown from one rat during wakefulness (W) and ketamine anaesthesia (K). Spectral exponents from all rats are also shown (*inset*). **c**, *Left*, time courses of mean PCI^*ST*^ in wakefulness, ketamine, and propofol (P) anaesthesia (shades represent SEM; horizontal lines indicate periods of statistical difference, *P* < 0.05). *Right*, PCI^*ST*^ in range 0.08-0.6 s is shown for each rat. Propofol data are the same as in Fig. 1. **d**, Number of principal components and **e**, average state transitions of EEG response are shown for all rats. **f**, Mean HF power (in range 0.08-0.18 s) and **g**, time of ITPC drop averaged across channels are shown for all animals during wakefulness and ketamine anaesthesia. **h**, Ratio between the number of electrodes (channels) with a late increase in HF power (after 0.08 s) and the total number of channels. **i**, Temporal differences between the onset of the late HF power and ITPC drop are shown.

We also used another general anaesthetic, the volatile sevoflurane, to test whether OFF periods combined with reduced ITPC and low PCI^*ST*^ were not specific to propofol. Sevoflurane anaesthesia in 10 rats produced results that were indistinguishable from what we observed with propofol (**Supplementary Fig. 4, 7, Video 6**).

### Functional connectivity and diversity of response were conserved during wakefulness and ketamine anaesthesia, but collapsed with propofol and sevoflurane

In principle, PCI estimates integration and differentiation in a neuronal network^1,11^. Thus, high PCI value should indicate a highly connected network with diversified connectivity pattern. In order to test this, we assessed the functional connectivity across cortical regions following electrical stimulation by computing the “inter-sites phase clustering” (ISPC)^23^ for each channel pair, in the frequency range 5-14 Hz, in 2 time windows of interest: during the OFF period (0.08-0.18 s) and afterwards, until the mean ITPC drop time in wakefulness, post OFF (0.18-0.3 s). We considered only the increase in ISPC above the baseline and computed the connectivity degree (CD) for each electrode, as the ratio between the number of significantly synchronized channel pairs over the total number of channels (**Fig. 5a**). During wakefulness, the averaged CD was sustained up to 0.3 s and was higher than during propofol anaesthesia in both time windows (**Fig. 5b**; 9 rats; OFF period, wakefulness: 0.79 ± 0.03, propofol: 0.40 ± 0.07; post OFF, wakefulness: 0.70 ± 0.04, propofol: 0.12 ± 0.02; Wilcoxon S-R test, for both windows, *P* = 0.004). Sevoflurane gave similar results (**Supplementary Fig. 8**; 9 rats). In contrast, ketamine anaesthesia induced a significant drop of CD compared to wakefulness only during the OFF period (**Fig. 5c;** 7 rats; wakefulness: 0.81 ± 0.04, ketamine: 0.57 ± 0.05; Wilcoxon S-R test, *P* = 0.016). After the OFF period, no significant difference was identified (wakefulness: 0.77 ± 0.04, ketamine: 0.62 ± 0.06; Wilcoxon S-R test, *P* = 0.078). Coherently, CD after the OFF period was higher with ketamine than with both propofol and sevoflurane, while no difference was detected between these latter conditions (Mann-Whitney test, ketamine vs propofol, *P* = 0.001; ketamine vs sevoflurane, *P* = 0.003; propofol vs sevoflurane, *P* = 0.093). We then identified a highly significant positive correlation between mean CD_(0.18-0.3s)_ across channels and PCI^*ST*^_(0.08-0.6s)_, thus revealing a connection between functional connectivity and perturbational complexity (**Fig. 5d**; linear fit, *P* = 2.138*10^−10^, R^2^ = 0.721). Not only the overall amount of connectivity differed between conditions; averaging CD_(0.18-0.3s)_ across channels organized in cortical regions (frontal: M2, M1; parietal: S1, PA, RS; occipital: V2, V1), revealed an uneven spatial distribution of CD during both wakefulness and ketamine anaesthesia (**Fig. 5e**; wakefulness, Friedman test, *P* = 0.004; ketamine, Friedman test, *P* = 0.018), indicating a peak of connectivity in the occipital region. The spatial distribution of CD was similar between wakefulness and ketamine anaesthesia (Friedman test, *P* = 0.555) and no relation with cortical areas was detected during both sevoflurane and propofol anaesthesia (sevoflurane, Friedman test, *P* = 0.097; propofol, Friedman test, *P* = 0.459). This suggested a similar degree of diversity in the evoked response among cortical areas during both wakefulness and ketamine conditions, which collapsed with sevoflurane and propofol.

**Fig. 5.**
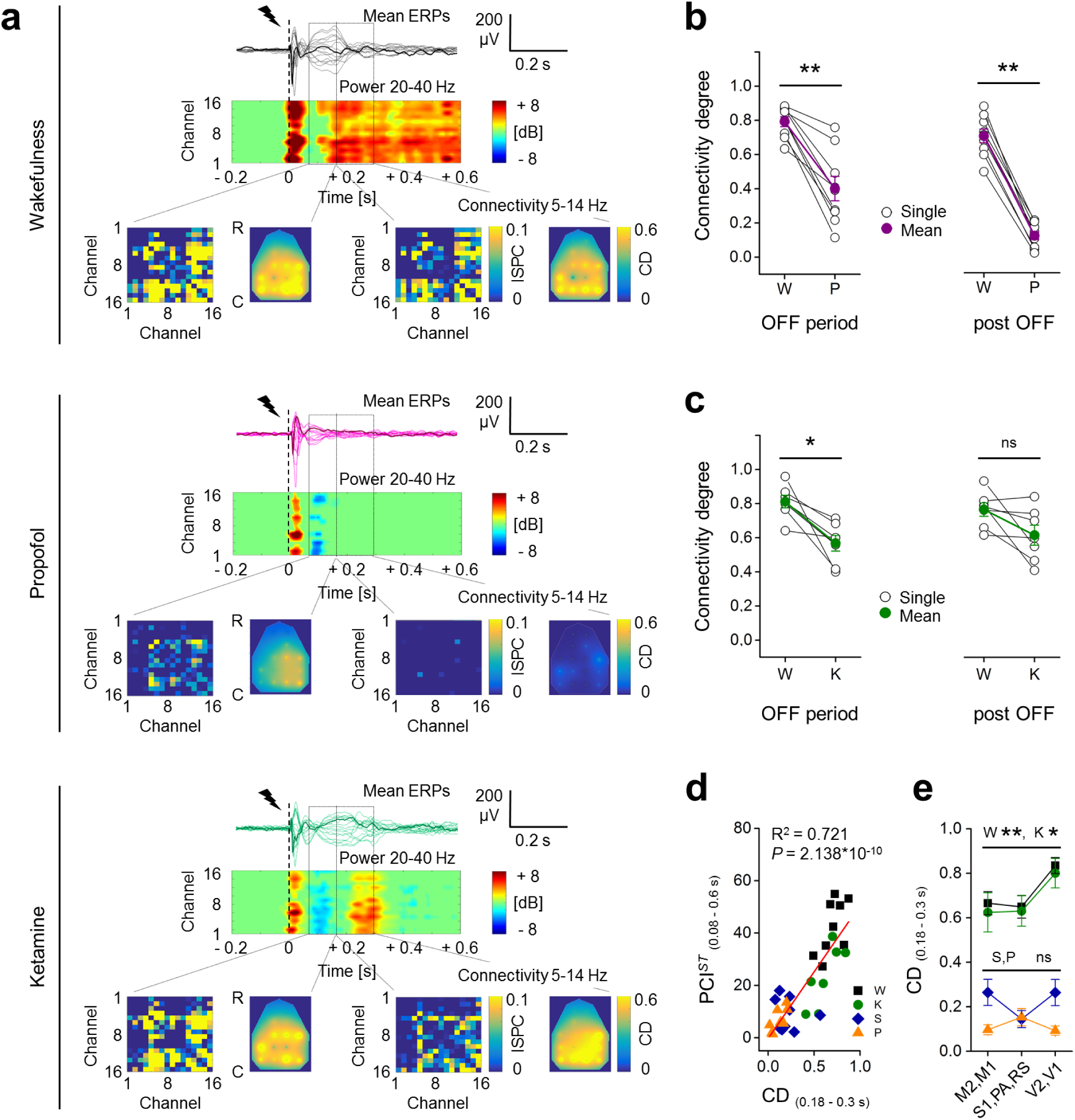
Functional cortical connectivity after perturbation was reduced during propofol or sevoflurane anaesthesia compared to wakefulness, while was conserved with ketamine. **a**, Superimposition of mean ERPs from all electrodes in response to single pulse stimulation (1 ms, 50 µA; dashed line) from the same rat during wakefulness (W) and propofol (P) and ketamine (K) anaesthesia (from *up* to *bottom*). One averaged ERP is in bold for clarity. Spectrograms of HF power are shown for each channel, below the butterfly plots. The bottom part of each inner panel reports increments in functional connectivity compared to baseline in two time windows: OFF period, 0.08-0.18 s, *left* and post OFF, 0.18-0.3 s, *right* (rectangles indicate the time windows). For each window, the connectivity matrix based on mean ISPC (5-14 Hz) is reported on the *left* and the topographical distribution (R-C: rostral-caudal) of CD for each channel is interpolated and shown on the *right*. **b-c**, Mean CD across channels in the OFF period, *left* and post OFF, *right*, from rats during wakefulness and propofol (**b**, n = 9), and ketamine (**c**, n =7) anaesthesia. **d**, Mean CD (range: 0.18-0.3 s) across channels from all animals and conditions are plotted against PCI^*ST*^ (range: 0.08-0.6 s) and linearly fitted (coefficient of determination R^2^ and *P* value are reported). **e**, Mean CD (range: 0.18-0.3 s) across rats and across channels organized in 3 cortical regions are shown for each condition. In **d** and **e** sevoflurane condition (S) is also reported (n = 9).

A complementary way to conceive the diversity in activity patterns induced by a stimulus is in relation to the specific site of stimulation. We took advantage of the variability, across rats, in the precise location of the stimulating electrode within M2, and tested for possible correlations with PCI^*ST*^. We did not find correlation with the position of the stimulating electrode along the rostro-caudal axis (in range: 3-5 mm from Bregma; **Supplementary Fig. 9**), but we detected a strong and positive correlation of PCI^*ST*^ with the location along the dorso-ventral axis (range: 0.1-0.83 mm from cortical surface, **Supplementary Fig. 9**). Specifically, during wakefulness PCI^*ST*^_(0.08-0.6s)_ positively correlated with the depth of the stimulation (**Fig. 6a-b**; linear fit, *P* = 0.003, R^2^ = 0.908) among 6 rats and a similar correlation was detected with ketamine (linear fit, *P* = 0.002, R^2^ = 0.922). By contrast, we did not find any significant correlation during both propofol and sevoflurane anaesthesia (**Fig. 6a-b;** linear fit, propofol: *P* = 0.387, sevoflurane: *P* = 0.082).

**Fig. 6.**
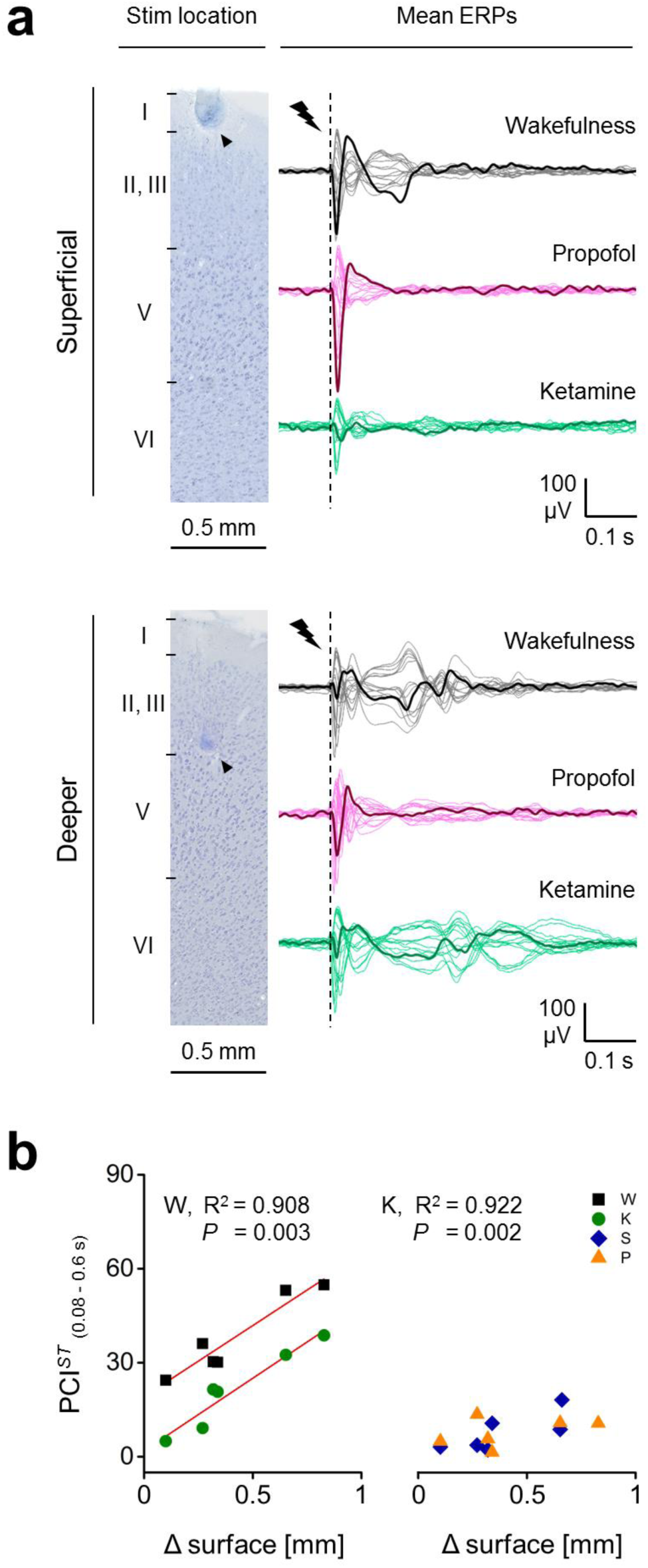
PCI^*ST*^ positively correlated with the depth of the stimulation site within M2 cortex during wakefulness and ketamine anaesthesia, but not with propofol or sevoflurane. **a**, *Left*, coronal cortical sections (Nissl staining) showing the location of the electrical stimulation in right M2 from one rat with the stimulating electrode positioned close to the cortical surface (*up*) and from another animal with the stimulating electrode deeper implanted in the cortex (*bottom*). Black arrowheads indicate the marks of one pole of the stimulating electrode. *Right*, Superimposition of mean ERPs from all recording electrodes in response to single pulse stimulation (1 ms, 50 µA; dashed line) from the same 2 rats shown on the left, during wakefulness (W) and propofol (P) and ketamine (K) anaesthesia. One averaged ERP from the same channel (S1) is in bold to highlight differences across conditions. **b**, Values of PCI^*ST*^ (in time range: 0.08-0.6 s) from 6 rats and for all conditions are plotted against the corresponding distances of the stimulating electrode tips from the cortical surface and linearly fitted (coefficient of determination R^2^ and *P* value are reported if *P* < 0.05). Strong positive correlations were identified for wakefulness and ketamine conditions with similar slopes (45.35 and 46.30 respectively), but not for propofol and sevoflurane (S) anaesthesia.

## DISCUSSION

We established a method for reliable, chronic recording of multichannel, epidural EEG in response to local cortical perturbation in rats (**Fig. 1**). For the first time in a non-human animal species *in vivo*, we quantified a variant of the Perturbational Complexity Index (PCI^*ST*^), a proposed measure of the brain’s capacity to sustain complex dynamics and conscious brain states^1,11^. We stimulated M2 because it is a highly integrated area in rodent cortex^28,29^, suitable for triggering widespread and differentiated responses, and it resembles human premotor cortex, often used for probing PCI^1,9^. We tested complexity during wakefulness and general anaesthesia induced by propofol, sevoflurane and ketamine, mimicking a previous study in humans^9^. The depth of anaesthesia was set at the minimal dose that abolished all motor responses to pain stimulation. At the same level of behavioural unresponsiveness, propofol and sevoflurane induced a spontaneous EEG activity characterized by high amplitude, low frequency waves, while ketamine anaesthesia yielded sustained fast, desynchronized EEG activity, resembling the awake state. In agreement with previous studies^30,31^, the spectral exponent of the relative periodogram (20-40 Hz) was reduced by propofol and sevoflurane, but with ketamine it remained similar to wakefulness (**Fig. 1, 4, Supplementary Fig. 7**). During wakefulness, the electrical stimulation produced a long-lasting and complex ERP with a sequence of reproducible voltage inversions in time and across cortical areas. The ERP had a peak power in the 8-25 Hz range, but reached up to 40 Hz and remained phase-locked for ∼0.3 s. Conversely, in propofol anaesthesia, the ERP was short-lasting, with few polarity changes, and after a brief HF response, ITPC dropped within ∼0.1 s (**Fig. 1-2**). Thus, PCI^*ST*^ was reduced from wakefulness to propofol anaesthesia in all rats (**Fig. 1, Supplementary Fig. 6**), in agreement with previous results from humans^9^.

Next, we investigated possible mechanisms underlying the reduced complexity. It has been hypothesized that cortical bistability may be important for preventing complex cortical dynamics^13,14,15^. Specifically, when the arousing input from the brainstem and thalamus is reduced or counteracted, such as during NREM sleep, general anaesthesia, or deafferentation, cortical circuits tend to become bi-stable and fall into synchronized down-states (neuronal hyperpolarization) following previous activations, or up-states^19,32,33,20,21^. At the network level, this phenomenon give rise to slow wave oscillations in the spontaneous EEG observed during NREM sleep and general anaesthesia, where periods of HF activity alternate with OFF-periods of HF suppression^19,20,17,18^. Neuronal down-states are thought to be generated mainly by activity-dependent K^+^ currents^34,35^ and/or synaptic fatigue and inhibition^36,37^ and it has been suggested that the same mechanisms may be triggered by the initial response to cortical stimulation. The “induced down-state” would then interrupt the deterministic and long-lasting sequence of complex neuronal interactions that are thought to give high PCI^13,14,15^. Indirect evidence for this mechanism was recently observed in humans during NREM sleep and UWS, when an OFF period was found to follow the initial response to cortical stimulation, and to correlate with the interruption of the phase-locked activation^12,10^. Similar results were obtained in cortical slices *in vitro*, with pharmacological reduction of bistability^24^. In agreement with these results, we detected a profound and widespread HF suppression, after the initial response, ∼0.08 s after stimulation, during propofol anaesthesia, in all animals (**Fig. 2, Suppementary. Fig. 5**). Importantly, by increasing the stimulus intensity during propofol anaesthesia, we were always able to obtain a similar cortical excitation as during wakefulness, but the OFF period was never abolished. Instead, both the magnitude and duration of the HF suppression increased linearly with the stimulus intensity (**Fig. 3**), suggesting that the OFF period likely reflected an adaptation-like “induced down-state”. This is in line with the idea that when bistablility dominates, a stronger activation leads to a more hyperpolarized state, and is compatible with results from slow waves in cortical slices, where an increased firing rate during the up-state was followed by longer and more hyperpolarized down-state^38^. Moreover, we found a positive temporal correlation between the end of the OFF period and the drop of ITPC (**Fig. 3**), with a strength which was surprisingly close to that observed in UWS and NREM sleep in humans^12,10^. Interestingly, the HF response was not simply abolished after the OFF period: stimulation-induced HF activations were still detectable later, even if they were sparse in space and time, and not phase-locked, more resembling a modulation of ongoing activity^22^. In contrast, during wakefulness, later HF activations were still phase-locked, sustained and widespread, indicating strong deterministic responses (**Fig. 2, 4**). Traces of HF suppression were detected also during wakefulness in response to stimulation, but these never interrupted durable phase-locked activations. Compared to propofol anaesthesia, the HF suppression during wakefulness was shallower, briefer and insensitive to variations in stimulus intensity (**Supplementary Fig. 5**), suggesting a different mechanistic origin. A similar effect during wakefulness was also previously observed in humans, from electrodes close to the stimulation site^39,12^, resembling a momentary increased recruitment of inhibition caused by vigorous electrical stimulation^39,12^.

Next, we examined the functional connectivity following the stimulation, within frequencies (5-14 Hz) that are implicated in long-range and feedback interactions across cortical areas^40,41^. We used ISPC as a measure of the consistency of phase-based connectivity across trials, to capture the spatial representation of the deterministic sequence of neuronal events that generates the complexity of the response in time. Consistently, we found a strong positive correlation between the derived CD and PCI^*ST*^(**Fig. 5**). Indeed, the response during wakefulness showed a high degree of connectivity with a peak in occipital areas, highlighting a certain diversity in the evoked neuronal activations. In contrast, propofol anaesthesia reduced cortical connectivity, and the spatial distribution of the remaining functional connections became more uniform^42^, resembling a reduction of both integration and differentiation of the neuronal response (**Fig. 5**). This is coherent with the known effect of propofol in suppressing long-range spontaneous network interactions^43^ and also with the reduced variability in the complexity of evoked responses to stimulations at different cortical depths, observed here (**Fig. 6**).

We next used the same approach during sevoflurane anaesthesia. Sevoflurane and propofol are known to share some molecular mechanisms, such as enhancement of GABA_A_ receptors^26,44^ and inhibition of voltage-gated Na^+^ channels^25,45^, but there are also important differences. Thus, sevoflurane opens 2-pore-domain (2P) K^+^ channels, unlike propofol^46^, and differently affects sensory processing^47^. Nevertheless, sevoflurane produced effects that were indistinguishable from those with propofol (**Supplementary Fig. 7-8, Fig. 5-6**). Taken together, these results support the hypothesis that the down-state may serve as a general mechanism for interrupting complex cortical dynamics. This mechanism seems indeed to be shared across anaesthetic drugs, animal species, and brain states, including NREM sleep^12^ and UWS^10^ in humans.

Nevertheless, our results with ketamine may seem to challenge this idea. Ketamine anaesthesia is known to produce a state of behavioural unresponsiveness with “dream-like”, vivid experiences^27^. A previous study in humans found no significant difference in PCI between wakefulness and ketamine anaesthesia^9^, but, signs of bistability have never been examined in this condition. Here we found that ERPs during ketamine showed mixed features: the initial brief response was rapidly followed (∼0.08 s later) by an OFF period, resembling propofol anaesthesia, but this did not prevent long-lasting deterministic activations in nearly half the animals, and the duration of the resulting phase-locked response was close to that of wakefulness (**Fig. 4**). After the OFF period, a later increase in HF activity was highly frequent, but neither consistently phase-locked and “deterministic”, like in wakefulness, nor consistently not phase-locked, like in propofol/sevoflurane conditions (**Fig. 4**). During ketamine anaesthesia, the time course of PCI^*ST*^ revealed similarities to wakefulness, but resulting in an overall reduction of complexity. However, PCI^*ST*^ from the onset of the OFF period was significantly higher than in propofol/sevoflurane anaesthesia, thus indicating an intermediate level of perturbational complexity with ketamine, regardless of the similar state of behavioural unresponsiveness (**Fig. 4**). Consistently, the functional connectivity across cortical regions was reduced compared to wakefulness during the OFF period, but recovered soon afterwards and a wakefulness-like pattern of CD across space was maintained, suggesting conserved integration and diversity (**Fig. 5**). Another important sign of a conserved differentiation during ketamine anaesthesia was the positive linear relation between PCI^*ST*^ and the depth of the site of stimulation within M2, which was also observed in wakefulness, with similar slopes (**Fig. 6**). The mixed results with ketamine may suggest some variability or unstable effects of this drug. In support to this idea, a recent study in humans found that the EEG activity spontaneously fluctuated between states of low and high complexity during ketamine anaesthesia, until awakening^48^. Certainly, since our experimental design was adopted to reproduce experiments in humans, we could only indirectly infer signs of down-states by detecting OFF periods in EEG. Indeed, HF suppression is a more direct indication of reduced neuronal activation relative to ongoing activity, and does not necessarily indicate neuronal hyperpolarization, which can be observed only by intracellular recordings. For example, a coordinated reduction in presynaptic release might also produce an OFF period in the EEG, if compared to the enhanced spontaneous activity that is typically induced by ketamine^49,50^, without implying hyperpolarization.

Although our data cannot yet resolve this issue, they illustrate the need for intracellular recordings to determine the role of bistability in the interruption of complex dynamics. Moreover, the novel correlation we identified between PCI^*ST*^ and the stimulus location within cortical depth represents a promising starting point for studying the role of specific cortical layers and micro circuitries in sustaining cortical complexity. In conclusion, we demonstrated that rodent cortical circuits can respond to focal stimulation with long-lasting sequences of deterministic complex interactions during wakefulness, which are disrupted during general anaesthesia, as previously shown in humans^9^. In agreement with previous results^12,24,10^, we provided indirect evidence for a connection between cortical bistability, interruption of deterministic responses, and disruption of cortical connectivity and complexity. Our results improve our understanding of the cortical dynamics that in humans have been associated with consciousness, and our method opens a range of future possibilities for more detailed, mechanistic investigations of brain states and their transitions.

## METHODS

### Animal model and ethical approval

All experimental and animal care procedures were carried out at the Institute of Basic Medical Sciences, University of Oslo (Norway) and were approved by the Norwegian Veterinary and Food Safety Authority (FOTS ID: 11812) in agreement with Norwegian law of animal handling. Experiments were carried out on adult male Sprague-Dawley rats (300-550 g, n = 12). All efforts have been made to avoid or minimize animals’ distress and pain during the entire course of experimentation. After implantation of chronic electrodes, rats were individually caged in enriched environments, with *ad libitum* access to food and water and were exposed to 12 h light-dark cycle at 23°C constant room temperature.

### Surgical procedure

All coordinates for electrodes implantation have been calculated from brain atlas of adult rats^1^ and are expressed referring to bregma position, X = medial-lateral axes (- left hemisphere, + right hemisphere), Y = rostral-caudal axes (- caudal to bregma, + rostral to bregma), Z = dorsal-ventral axes. Event related potentials (ERPs) were induced by electrical stimulations delivered by a custom made bipolar electrode chronically implanted in right secondary motor cortex (M2). The stimulating electrode was composed by 2 tungsten wires of 50 *μ*m caliber, PTFE insulated (Advent, Oxford, England), connected to gold pins (Mouser electronics, Upplands-Väsby, Sweeden) at one end. The other ending tips of the wires were cut in ∼45°, adjusted at the same horizontal plane and fixed in a parallel position, separated by ∼500 *μ*m. The resulting bipolar electrode was inserted perpendicularly to the cortical surface, along the coronal plane and the coordinates (in mm) for implantation were calculated as X = + 1.2 left wire / + 1.7 right wire; Y = + 3.7; Z = + 1.9. ERPs were recorded by 16 custom made electrodes plus one ground (GND), consisting of stainless steel screws (1.2 mm caliber; Centrostyle, Vedano Olona, Italy) soldered to a gold pin via a conductive insulated wire (Omnetic, Minneapolis, USA). The screws were in contact with the dura by penetrating the skull for recording epidural EEG. Efforts were made for not exceeding the internal surface of the skull with the flat base of the screws. Recording electrodes covered most of cortical areas, were organized in a grid symmetric along the sagittal suture and were placed at the following coordinates (in mm): X = ± 1.5, Y = + 5 (M2); X = ± 1.5, Y = + 2 (M2); X = ± 1.5, Y = - 1 (M1); X = ± 4.5, Y = - 1 (S1); X = ± 1.5, Y = - 4 (RS); X = ± 4.5, Y = - 4 (PA); X = ± 1.5, Y = - 7 (V2); X = ± 4.5, Y = - 7 (V1); X = 0, Y = - 10 (cerebellum, GND). Impedances of all electrodes were measured *in situ* at 1 kHz at the beginning of each recording session, for each rat. The averaged impedance (mean ± SEM) across all sessions, channels and animals was 0.17 ± 0.03 MΩ for stimulating electrodes and 7.12 ± 0.42 kΩ for recording electrodes.

For electrode implantation, rats were deeply anaesthetized in an induction chamber (sevoflurane 5 %, Baxter, Oslo, Norway) until loss of righting reflex. Rats were then connected through a gas mask to an anaesthesia delivery system (SomnoSuite, Kent Scientific, Torrington, CT, USA). Absence of response to deep tail pain stimulations was verified, then rats were fixed in a stereotaxic frame (TSE systems, Bad Homburg, Germany). During surgical procedure, deep general anaesthesia (sevoflurane 2.5 - 5 %) was constantly delivered in oxygen concentrated (O2 > 85 %) humidified room air (rate 0.5 l/min), body temperature was maintained at 36.5 - 37.5 °C by a heating pad (DC Temperature Controller, FHC, Bowdoin, ME, USA) and lactated ringer’s solution (Fresenius Kabi, Halden, Norway) was administered intravenously with a syringe pump (sp220i, WPI, Hertfordshire, UK) through the tail vein (rate 3.3 ml/kg/h) to maintain body fluids. Rats received subcutaneous injections of butorphanol (2 mg/kg MSD, Boxmeer, Nederland) and dexamethasone (0.2 mg/kg, MSD, Boxmeer, Nederland) to ensure analgesia and prevent inflammatory response, and standard sterile procedures were used throughout the surgical operation^2^. Holes were drilled through the exposed skull at the desired coordinates, electrodes were positioned and two machine screws were also upside-down mounted over the caudal part of the skull for subsequent head restriction. Dental acrylic cement (Meliodent, Kulzer, Hanau, Germany) was applied over the entire exposed skull, sealing the wound margins and securing electrodes and screws in place. After surgery, buprenorphine (0.1 mg/kg, Indivior, Slough, Uk) and meloxicam (1 mg/kg, Norbrook, Uppsala, Sweeden) were subcutaneously injected. For the following 3-4 days rats were checked for possible signs of distress, infection or for damages to electrode implantation. Buprenorphine (0.1-0.05 mg/kg) and dexamethasone (0.2-0.1 mg/kg) were also subcutaneously injected once a day.

### Experimental procedures

After recovery, rats were gradually habituated to body and head restriction and acclimatized to a custom recording setup in 3 consecutive days. Rats were anesthetized (brief exposure to sevoflurane 5 %), quickly placed in a custom made bag for restricting body movements and introduced into an elevated acrylic tube. Only the head and the tail were left outside the acrylic tube. After awakening, rats were gradually kept in the same condition of body restriction for longer times (first day 30 min, second day 45 min, third day 60 min). Head restriction by a clamped head-bar connected to the machine screws was gradually introduced from the second day of habituation and drops of mango juice were given as reward. After 1 week from surgery, rats underwent several electrophysiological stimulation/recording sessions at first during wakefulness followed by general anaesthesia. Recording sessions were interleaved with a resting period of at least 48 hours. Three different general anaesthetics were tested (sevoflurane, propofol and ketamine) in randomized different days and most of the rats were exposed to all anaesthetic agents. At the end of the last recording session, during general anaesthesia, an electrolytic lesion was performed by applying 30 µA for 30 s to the poles of the stimulating electrode. Rats were then sacrificed with a lethal dose of pentobarbital (140 mg/kg, i.v., Virbac, Carros, France) and after the suppression of corneal reflex, were intracardially perfused with PBS (phosphate buffer solution with heparin 5000 IU/l, LEO-Pharma, Lysaker, Norway) and 4 % paraformaldehyde (Sigma-Aldrich, St. Louis, MO, USA) in PBS at 4 °C for tissue fixation. Brains were then extracted and processed for histological staining.

During the recording sessions rats were placed in the same setup and conditions of body and head restriction used for habituation and the tail vein was also cannulated (26 G catheter) for allowing i.v. infusion of propofol or ketamine, when needed. The stimulating electrode was connected to an isolated current stimulator (Isolator HG203, High Medical, London Uk) triggered by a voltage pulse generator (2100, A-M System, Washington DC, USA). Recording electrodes were connected to a 16-channel unipolar amplifier board with common reference shorted to ground (RHD2132, Intan Technologies, Los Angeles, CA, USA), controlled by Open Ephys system^3^, and the epidural EEG signal was acquired and digitized at 10 or 30 kHz, 16-bit resolution.

Stimulation/recording was carried out in darkness after ∼45 min of acclimatization in the setup. Depending on the randomized recording session, rats were exposed to several electrical monophasic current pulses (duration 1 ms) of 50 *μ*A or of various intensities (40, 60, 80 and 100 *μ*A, organized in randomized blocks) delivered at 0.1 Hz during wakefulness. Stimulations were repeated during subsequent general anaesthesia within the same animal. General anaesthesia was initially induced either by the exposure to sevoflurane 5 %, or by the i.v. bolus injection of propofol 10 mg/kg (B-Braun, Melsungen, Germany) or ketamine 30 mg/kg (Vetoquinol, Ittigen, Swiss). General anaesthesia was then maintained at the initial constant dosage of either sevoflurane 2.5 % (by a gas mask, in humidified air, O2 > 85 %, at 0.5 l/min), propofol 1 mg/kg/min or ketamine 1.75 mg/kg/min (by a syringe pump). The gas mask was also positioned during conditions of propofol and ketamine anaesthesia in order to deliver oxygen (as during the exposure to sevoflurane). Subcutaneous injection of glycopyrrolate 0.01 mg/kg (Meda, Asker, Norway) was also performed during ketamine anaesthesia to reduce the induced increase in salivation^4^. Lubrithal eye ointment (Dechra, Hadnall, UK) was applied to keep eyes moist and body temperature was kept at 36.5 - 37.5 °C by a heating blanket system (Homeothermic Monitor, Harvard Apparatus, Holliston, MA, USA). The initial dosage was the minimal necessary for maintaining loss of righting reflex in 100 % of the tested rats^5,6^. After 10 min from induction, rats were checked for reaction to pain stimulations (pinching the tail by a forceps) and the concentration or infusion rates were stepwise increased by adding 4 % of the initial dosage until any reflex was absent (3 min passed between each increment). The resulting averaged experimental dosages across rats and sessions were (mean ± SEM): sevoflurane 2.58 ± 0.03 %, propofol 1.06 ± 0.02 mg/kg/min and ketamine 1.83 ± 0.03 mg/kg/min. EEG recordings that showed sustained burst suppression induced by deep general anaesthesia have been excluded from analysis.

A subset of animals were exposed to a final stimulation/recording session with the purpose to detect putative movements of whiskers induced by electrical stimulation of M2. Whiskers of the left mystacial pad (contralateral to the site of stimulation) were clipped, except for the most caudal vibrissa of the third row of the mystacial pad (C1), which was tracked by a high speed camera (500 f/s, Blackfly S Mono 0.4 MP USB3, FLIR System, Wilsonville, OR, USA). Experimental area was constantly illuminated only by dim red light (wavelength 655 nm, LED, Quadica, Lethbridge, Canada), which is not detectable by rat retina^7,8^, but allowed video recording. At the same time, a second LED 655 nm was triggered by the stimulation system, signaling the onset of current pulses to the camera. During wakefulness, head and body restrained rats were then exposed to several monophasic current pulses of 1 ms at either 50 *μ*A or 100 *μ*A, or to train stimulations of duration 0.3 s composed by 11 single monophasic current pulses of 1 ms and 50 *μ*A (33 Hz) as positive control. Different stimulations were grouped in randomized blocks and delivered at a rate of 0.2 Hz. In order to avoid saturation artifacts in the amplification system during train stimulation, the epidural EEG activity was recorded by a bipolar amplifier board (RHD2216, Intan Technologies, Los Angeles, CA, USA) and the frontal-occipital (M2-V2) derivation, ipsilateral to the stimulation, was adopted for only this session. At the end of stimulation/recording, rats were deeply anaesthetized (sevoflurane 5 %), scarified with a lethal dose of pentobarbital (140 mg/kg, i.v) and processed as described above.

### Analysis of epidural EEG signal

Analysis of electrophysiological data was performed in MATLAB2016a (Math Works, Natick, Massachusetts, USA) and Origin 9.1 (OriginLab, Northampton, Massachusetts, USA). Raw epidural EEG recordings were visually inspected to remove channels containing noise artifacts or having impedance > 1 MΩ. EEG signals from all electrodes were re-referenced to the common average across channels for analyzing ERPs, while a bipolar frontal-occipital derivation (M2-V2 right) was chosen for analyzing spontaneous activity. Stimulus artifacts were removed and signal was spline interpolated in a time window from 0 to 0.005 s from stimulus onset. EEG signal was band pass filtered from 0.5 to 80 Hz (butterworth filter, 3^th^ order) and down-sampled to 500 Hz. ERP epochs from −5 to 5 s centered at the stimulus onset (0 s) were then extracted for each channel. All epochs were offset corrected by subtracting the average voltage of their respective baseline (from −1 to 0 s). Trials with high voltage artifacts in their baseline were removed. Threshold for rejection was set to the averaged root mean square of baseline (rms, from −1 to 0 s) across trials + 3 standard deviations. The first n = 90 consecutive trials of preprocessed signal were used for analysis of evoked responses to electrical perturbations of 50 µA and spontaneous EEG activity was quantified from baseline epochs of 4 seconds (from −5 to −1 s) from the corresponding trials (with the exception of one animal in one wakefulness condition, for which 80 trials were used in analysis instead). Whereas n = 56 consecutive ERPs were used for analysis of evoked responses to current simulations of increasing intensities.

In order to quantify the different states of brain activity induced by general anaesthetics in relation to wakefulness, fast Fourier transform (FFT) was performed on n = 90 epochs of spontaneous EEG activity and normalized by the number of samples N. The squared modules of the normalized FFTs were computed and the resulting power spectrums were averaged across trials. The 1/f relation of the averaged periodogram was then linearly fitted in Log-Log coordinates, in the frequency range from 20 to 40 Hz. The slope of the obtained linear function was considered to be the spectral exponent of the 1/f function and was used to quantify the (re)distribution of frequency powers in the spontaneous EEG activity^9^.

Several features of the ERPs were evaluated. The actual excitation of cortical tissue induced by electrical perturbation was quantified by the rms amplitude of the first deflection of the mean ERP (0.006 - 0.05 s from stimulus onset) from each electrode and then averaged across channels. The same quantification was also performed on a later deflection of evoked potential around 0.2 s (from 0.175 to 0.225 s).

Morlet wavelet convolution was performed on each trial for all channels to extract both spectral powers and phases of ERPs. A family of 40 wavelets (3 cycles) linearly spanning from 1 to 40 Hz was adopted. Powers from each trial and frequency were normalized over the averaged power across trials in the baseline time window from −0.5 to −0.2 s for each respective frequency. The relative powers were then averaged across trials for each channel and frequency and converted in dB. In order to identify only the significant variations of power with respect to the baseline, bootstrap statistic (500 trial-based permutations) was performed. One positive threshold and one negative threshold (both, *α* = 0.05) were identified for each frequency and channel, based on the obtained distribution of the maximum and minimum dB values in the baseline time window. All values of relative power equal to or in between the corresponding thresholds were then set to 0 and only the significant mean relative powers across trials were considered for the analysis. A global approximation of the spectral content of the ERP was quantified by averaging the relative powers in frequency bands: δ (1 to 4 Hz), θ (5 to 7 Hz), α (8 to 14 Hz), β (15 to 25 Hz) and γ (26 to 40 Hz), then across time (from 0 to 0.5 s) and across channels. The presence of a reduced power in high frequency range (HF > 20 Hz) after stimulation, relative to baseline (HF power < 0 dB) was considered to be an indirect sign of neuronal silence and may indicate a neuronal down-state of cortical neurons in response to stimulation^10,11,12,13^. Since the period of neuronal silence is here inferred from extracellular recordings rather than direct measures of membrane potential, the terms “OFF period” or “HF suppression” are preferred^12,13^. Relative powers of ERPs were averaged in the frequency range of 20-40 Hz, and starting and ending time points of OFF periods were identified in a time window from 0 to 0.3 s as first downward and last upward zero crossing of the resulting dB signal. Minimum dB peak in the same time window was also detected, and the HF power was quantified by averaging the relative power in the frequency range of 20-40 Hz, from 0.08 to 0.18 s. The starting point of the time window used to quantify HF power was empirically determined and was the mean starting point of OFF period across anaesthetic conditions. Potential later increments of HF power (> 0 dB) were detected in a time window from 0.08 to 0.8 s. All the acquired values were then averaged across channels, obtaining approximations for each rat.

In order to quantify the deterministic effect that the electrical stimulation had on the cascade of neuronal events generating the EEG response, we measured the duration of increased phase-locking of subsequent ERPs compared to baseline^12,13^. Phases of all time-frequency-trial-channel points were extracted from wavelet convolution and phase-locking at each frequency and time point was computed as “inter-trial phase clustering” (ITPC)^14^ with the following formula:

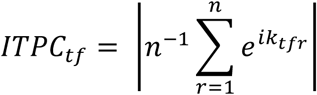

Where *n* is the number of trials and *e*^*ik*^ is the complex polar representation of the phase *k* from the trial *r* at time-frequency point *tf*. ITPC is an adimensional measure and can assume values from 0 (no phase locking) to 1 (maximal phase locking). Trial-based bootstrap statistic (500 permutations) was performed in order to conserve only the statistically significant increments of ITPC with respect to baseline (from −0.5 to −0.2 s). A threshold (*α* = 0.01) was identified for each frequency and channel based on the obtained distribution of the maximum ITPC values in the baseline time window, and all ITPC values found to be equal to or lower than the respective threshold were set to 0. Significant ITPC values were averaged across a broad band frequency range between 8 to 40 Hz^12,13^ and the “ITPC drop” was defined as the time point of the last ITPC value higher than 0 in a time window from 0 to 0.8 s from the stimulus onset. ITPC drop time values from all channels were then averaged to obtain an approximation for each rat.

A similar phase-based approach was adopted to quantify increments in functional connectivity across channels following the perturbation compared to baseline. For each trial, phase differences across channels at each frequency and time point were computed and the clustering over trials of resulting phase differences was defined as “inter-site phase clustering” (ISPC)^14^ and calculated with the following formula:

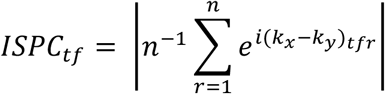

Where *n* is the number of trials and 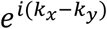 is the complex polar representation of the difference between the phases *k*_*x*_ and *k*_*y*_ from the channels *x* and *y*, for the trial *r* at time-frequency point *tf.* Therefore, ISPC represents the consistency of the phase configuration between the activity from two channels across trials, at each time-frequency point. ISPC of each channel pair and frequency is then baseline corrected by subtracting the corresponding average value in the time window from −0.5 to −0.2 s and bootstrap statistic (500 trial-based permutations) was performed to maintain only the statistically significant ISPC values. A threshold (*α* = 0.01) was identified for each frequency and channel pair based on the obtained distribution of the maximum ISPC values in the baseline (from −0.5 to −0.2 s), and all ISPC values found to be equal to or lower than the respective threshold were set to 0. Moreover, all ISPC scores that could be explained by volume conduction (clustering of phase difference around 0 or π) were excluded from the analysis and set to 0. Resulting ISPC values from each channel pair were then averaged in the frequency range 5-14 Hz for in two time windows, during the OFF period (from 0.08 to 0.18 s) and immediately later (from 0.18 to 0.3 s). The resulting ISPC scores > 0 represented significant functional connections between channel pairs and the “connectivity degree” (CD) was defined as the number of significant functional connections for each electrode, normalized over the number of channels minus one (to exclude autocorrelation). CDs from all channels were then averaged for each time window, obtaining global approximations for each rat. Rats with more than 2 bad channels (broken or noisy electrodes) were excluded from the connectivity analysis.

The spatiotemporal complexity of EEG responses to electrical stimulations was calculated using the recently introduced “Perturbational Complexity Index state-transition” (PCI^*ST*^)^15^, which quantifies the number of state transitions present in the principal components of the signal’s response. Briefly, the principal components accounting for 99 % of the variance present in the response are obtained through singular value decomposition and then selected based on a minimum signal-to-noise ratio (SNR_min_). Then, for each component the number of significant state transition (ΔNST) – a metric derived from recurrent quantification analysis (RQA)^16^ is computed. PCI^*ST*^ is the product between the number of principal components surviving dimensionality reduction and the average number of state transitions across components^15^. Hence, PCI^*ST*^ is high when a response displays multiple linearly independent components (spatial differentiation), each contributing with significant amounts of state transitions (temporal complexity). In order to minimize the amount of baseline-like oscillations (noise) that contributed to the PCI^*ST*^ value, SNR_min_ was chosen using a bootstrap procedure in the following way: for each signal, PCI^*ST*^ was calculated on 16 surrogates for which complexity should be zero, generated using two random non-response 0.5 s segments (t < 0 s or t > 1.5 s, where 0 s is the stimulation onset) as baseline and response. SNR_min_ was then set to 1.8 for all analysis, the smallest value for which the median PCI^*ST*^ across all surrogates was zero. The baseline and response window were defined as (from −0.5 to −0.005 s) and (from 0 to 0.6 s). To assess how the complexity of the EEG responses varied in time, PCI^*ST*^ was calculated in shorter 0.1 s sliding windows from stimulus onset (0.02 s overlap) and in the time window from 0.08 to 0.6 s. PCI^*ST*^ was computed using the code available at <u>github.com/renzocom/PCIst</u> and further parameters were set as previously reported^15^.

### Analysis of whisker tracking

The video recording of the whisker movements in response to electrical stimulation was initially analyzed in Bonsai software^17^. The centroid of the whisker was tracked offline, and the relative space coordinates in the Cartesian plane were extracted for each frame and imported in Matlab. Whisker positions were converted into degrees in the polar plane, obtaining angular oscillations in time that were analyzed similarly to the voltage signal. From the continuous signal, n = 21 consecutive motor responses centered on the stimulus onset (0 s) were extracted (from −2.5 to 2.5 s). Each motor trial was offset corrected by subtracting the respective mean angle of the baseline (time window from −1 to 0 s) and all the analyses were performed at the level of single trials for each rat. The magnitude of the whisker oscillation in response to the stimulus was quantified by the rms amplitude in a time window of 0.25 s following the stimulus offset and compared with the rms of the baseline (from −0.5 to −0.25 s). The mean spectral power of the whisker oscillation was also computed. A 3-cycle Morlet wavelet convolution was performed with a family of 100 wavelets spanning linearly from 1 to 100 Hz. The powers of each frequency from all trials were extracted and normalized over the corresponding mean power across trials in the baseline (from −0.8 to −0.3 s). The relative powers of each frequency were then averaged over trials and converted into dB. Bootstrap statistic (500 trial-based permutations, thresholds *α* = 0.05) was performed and the non-significant angle variations with respect to the baseline were set to 0. The resulting relative powers were then averaged in a broad band frequency range (from 5 to 100 Hz), in the first 0.25 s after the stimulus offset.

### Statistics

All results are expressed as mean ± SEM (average across rats), error bars represent SEM and nonparametric statistics were adopted. In a repeated measure design with dependent variables having more than 2 levels, principal effect of the variable was tested with Friedmans test. Group comparisons in repeated measures design were tested with Wilcoxon signed-rank (S-R) test, otherwise Mann-Whitney test was adopted. Linear fitting was performed with the least-square method and error bars were used as weights when averages across rats were fitted. To evaluate correlations and goodness of fit, the coefficient of determination R^2^ was computed and t test was performed to test the null hypothesis of slope = 0, establishing a *P* value. Gaussian v-test was used to test volume conduction in connectivity analysis, and therefore if phase differences were significantly different from 0 or π^14^. All statistics are two-tailed. The statistical significance in figures are represented as follows: *P* < 0.05 *, *P* < 0.01 **, *P* < 0.001 ***, *P* ≥ 0.05 ns (not significant).

### Histological staining

After fixation, brains were exposed to increasing concentrations of sucrose (10, 20, 30 %) in PBS solutions at 4 °C for 4 days. Brains were quickly frozen in sucrose 30 % in PBS and sliced in coronal sections of 50 µm thickness with a HM 450 sliding microtome (ThermoFisher Scientific, Oslo, Norway). Coronal sections were then prepared for Nissl staining. Sections were first dehydrated in increasing concentration of ethanol (70, 95, 100 %) and immersed in xylene (VWR, Oslo, Norway) for removal of fat. Slices were rehydrated with decreasing concentration of ethanol (100, 95, 70, 50 %) and stained with Cresyl echt violet solution (incubation at 60°C for 13 min, Abcam, Cambridge, UK). Sections were then rinsed in H_2_O_dd_, dehydrated in ethanol and then mounted and secured with coverslip on microscope slides. Brain sections were scanned at 5x with an AxioScanZ1 slidescanning microscope (Carl Zeiss, Neubeuern, Germany) and estimation of electrode positions was conducted using ZEN imaging light software (Carl Zeiss, Oberkochen, Germany).

## ADDITIONAL INFORMATION

### Authors’ Contribution

JFS, AA conceptualized the experiments; AA designed the experiments; AA, ST performed experiments and collected data; AA analyzed data; RC, AGC performed PCI^*ST*^ analysis; ST performed Nissl staining; AA, JFS, AGC, RC wrote the manuscript; all authors participated in the interpretation of results and revision of the manuscript, and approved the final version of the manuscript.

## Acknowledgments

We would like to thank Per M. Knutsen for help in preparation of the experimental set-up and Charlotte Boccara for advice regarding the writing. We are especially grateful to Marcello Massimini, Andrea Pigorini, Matteo Fecchio and Simone Russo for their valuable comments, suggestions, and support along the way.

## ARTICLE REFERENCES

1. Casali, A. G. et al. A theoretically based index of consciousness independent of sensory processing and behavior. Sci. Transl. Med. 5, 1–10 (2013).

2. Dehaene, S. Consciousness and the brain: Deciphering how the brain codes our thoughts. (Penguin, 2014).

3. Koch, C., Massimini, M., Boly, M. & Tononi, G. Neural correlates of consciousness: progress and problems. Nat. Rev. Neurosci. 17, 307–21 (2016).

4. King, J. R. et al. Information sharing in the brain indexes consciousness in noncommunicative patients. Curr. Biol. 23, 1914–1919 (2013).

5. Schartner, M. et al. Complexity of multi-dimensional spontaneous EEG decreases during propofol induced general anaesthesia. PLoS One 10, 1–21 (2015).

6. Demertzi, A. et al. Human consciousness is supported by dynamic complex patterns of brain signal coordination. Sci. Adv. 5, 1–12 (2019).

7. Luppi, A. I. et al. Consciousness-specific dynamic interactions of brain integration and functional diversity. Nat. Commun. 10, (2019).

8. Casarotto, S. et al. Stratification of unresponsive patients by an independently validated index of brain complexity. Ann. Neurol. 80, 718–729 (2016).

9. Sarasso, S. et al. Consciousness and complexity during unresponsiveness induced by propofol, xenon, and ketamine. Curr. Biol. 25, 3099–3105 (2015).

10. Rosanova, M. et al. Sleep-like cortical OFF-periods disrupt causality and complexity in the brain of unresponsive wakefulness syndrome patients. Nat. Commun. 9, 1–10 (2018).

11. Comolatti, R. et al. A fast and general method to empirically estimate the complexity of brain responses to transcranial and intracranial stimulations. Brain Stimul. 12, 1280–1289 (2019).

12. Pigorini, A. et al. Bistability breaks-off deterministic responses to intracortical stimulation during non-REM sleep. Neuroimage 112, 105–113 (2015).

13. Tononi, G., Boly, M., Massimini, M. & Koch, C. Integrated information theory: from consciousness to its physical substrate. Nat Rev Neurosci 17, 450–461 (2016).

14. Sanchez-Vives, M. V., Massimini, M. & Mattia, M. Shaping the default activity pattern of the cortical network. Neuron 94, 993–1001 (2017).

15. Storm, J. F. et al. Consciousness regained: disentangling mechanisms, brain systems, and behavioral responses. J. Neurosci. 37, 10882–10893 (2017).

16. Brecht, M. et al. Organization of rat vibrissa motor cortex and adjacent areas according to cytoarchitectonics, microstimulation, and intracellular stimulation of identified cells. J. Comp. Neurol. 479, 360–73 (2004).

17. Steriade, M., Contreras, D., Amzica, F. & Timofeev, I. Synchronization of fast (30-40 Hz) spontaneous oscillations in intrathalamic and thalamocortical networks. J. Neurosci. 16, 2788–2808 (1996).

18. Mukovski, M., Chauvette, S., Timofeev, I. & Volgushev, M. Detection of active and silent states in neocortical neurons from the field potential signal during slow-wave sleep. Cereb. Cortex 17, 400–414 (2007).

19. Steriade, M., Nuñez, A. & Amzica, F. A novel slow (< 1 Hz) oscillation of neocortical neurons in vivo: depolarizing and hyperpolarizing components. J. Neurosci. 13, 3252–3265 (1993).

20. Steriade, M., Timofeev, I. & Grenier, F. Natural waking and sleep states: a view from inside neocortical neurons. J. Neurophysiol. 85, 1969–1985 (2001).

21. Volgushev, M., Chauvette, S., Mukovski, M. & Timofeev, I. Precise long-range synchronization of activity and silence in neocortical neurons during slow-wave sleep. J. Neurosci. 26, 5665–5672 (2006).

22. David, O., Kilner, J. M. & Friston, K. J. Mechanisms of evoked and induced responses in MEG/EEG. Neuroimage 31, 1580–1591 (2006).

23. Cohen, M. X. Analyzing neural time series data: theory and practice. (MIT Press, 2014).

24. D’Andola, M. et al. Bistability, causality, and complexity in cortical networks: An in vitro perturbational study. Cereb. Cortex 28, 2233–2242 (2018).

25. Ouyang, W., Wang, G. & Hemmings, H. C. Isoflurane and propofol inhibit voltage-gated sodium channels in isolated rat neurohypophysial nerve terminals. Mol. Pharmacol. 64, 373–381 (2003).

26. Bieda, M. C. & MacIver, M. B. Major role for tonic GABAA conductances in anesthetic suppression of intrinsic neuronal excitability. J. Neurophysiol. 92, 1658–67 (2004).

27. Collier, B. B. Ketamine and the conscious mind. Anaesthesia 27, 120–134 (1972).

28. Zingg, B. et al. Neural networks of the mouse neocortex. Cell 156, 1096–1111 (2014).

29. Barthas, F. & Kwan, A. C. Secondary motor cortex: where ‘sensory’ meets ‘motor’ in the rodent frontal cortex. Trends Neurosci. 40, 181–193 (2017).

30. Colombo, M. A. et al. The spectral exponent of the resting EEG indexes the presence of consciousness during unresponsiveness induced by propofol, xenon, and ketamine. Neuroimage 189, 631–644 (2019).

31. Lendner, J. D. et al. An Electrophysiological Marker of Arousal Level in Humans. bioRxiv 1–46 (2019). doi:10.1101/625210

32. Timofeev, I., Grenier, F., Bazhenov, M., Sejnowski, T. J. & Steriade, M. Origin of slow cortical oscillations in deafferented cortical slab. Cereb. Cortex 10, 1185–1199 (2000).

33. Sanchez-Vives, M. V & McCormick, D. a. Cellular and network mechanisms of rhythmic recurrent activity in neocortex. Nature neuroscience 3, 1027–1034 (2000).

34. Timofeev, I., Grenier, F. & Steriade, M. Disfacilitation and active inhibition in the neocortex during the natural sleep-wake cycle: An intracellular study. Proc. Natl. Acad. Sci. 98, 1924–1929 (2001).

35. Compte, A., Sanchez-Vives, M. V., McCormick, D. A. & Wang, X.-J. Cellular and Network Mechanisms of Slow Oscillatory Activity (<1 Hz) and Wave Propagations in a Cortical Network Model. J. Neurophysiol. 89, 2707–2725 (2003).

36. Esser, S. K., Hill, S. L. & Tononi, G. Sleep homeostasis and cortical synchronization: I. Modeling the effects of synaptic strength on sleep slow waves. Sleep 30, 1617–1630 (2007).

37. Funk, C. M. et al. Role of Somatostatin-Positive Cortical Interneurons in the Generation of Sleep Slow Waves. J. Neurosci. 37, 9132–9148 (2017).

38. Sanchez-Vives, M. V. et al. Inhibitory Modulation of Cortical Up States. J. Neurophysiol. 104, 1314–1324 (2010).

39. Borchers, S., Himmelbach, M., Logothetis, N. & Karnath, H.-O. Direct electrical stimulation of human cortex — the gold standard for mapping brain functions? Nat. Rev. Neurosci. 13, 63–70 (2012).

40. Von Stein, A. & Sarnthein, J. Different frequencies for different scales of cortical integration: From local gamma to long range alpha/theta synchronization. Int. J. Psychophysiol. 38, 301–313 (2000).

41. van Kerkoerle, T. et al. Alpha and gamma oscillations characterize feedback and feedforward processing in monkey visual cortex. Proc. Natl. Acad. Sci. 111, 14332–14341 (2014).

42. Juel, B. E., Romundstad, L., Kolstad, F., Storm, J. F. & Larsson, P. G. Distinguishing anesthetized from awake state in patients: a new approach using one second segments of raw EEG. Front. Hum. Neurosci. 12, 1–14 (2018).

43. Lewis, L. D. et al. Rapid fragmentation of neuronal networks at the onset of propofol-induced unconsciousness. Proc. Natl. Acad. Sci. 109, E3377–E3386 (2012).

44. Sebel, L. E., Richardson, J. E., Singh, S. P., Bell, S. V & Jenkins, A. Additive effects of sevoflurane and propofol on gamma-aminobutyric acid receptor function. Anesthesiology 104, 1176–1183 (2006).

45. Ouyang, W., Herold, K. F. & Hemmings, H. C. Comparative effects of halogenated inhaled anesthetics on voltage-gated Na+ channel function. Anesthesiology 110, 582–590 (2009).

46. Putzke, C. et al. Differential effects of volatile and intravenous anesthetics on the activity of human TASK-1. Am. J. Physiol. Cell Physiol. 293, C1319–C1326 (2007).

47. Arena, A. et al. Linear transformation of the encoding mechanism for light intensity underlies the paradoxical enhancement of cortical visual responses by sevoflurane. J. Physiol. 595, 321–339 (2017).

48. Li, D. & Mashour, G. A. Cortical dynamics during psychedelic and anesthetized states induced by ketamine. Neuroimage 196, 32–40 (2019).

49. Brown, E. N., Lydic, R. & Schiff, N. D. General Anesthesia, Sleep, and Coma. N. Engl. J. Med. 363, 2638–2650 (2010).

50. Ferro, M. et al. Functional mapping of brain synapses by the enriching activity-marker SynaptoZip. Nat. Commun. 8, (2017).

## METHOD REFERENCES

1. Paxinos, G. & Watson, C. The rat brain in stereotaxic coordinates. (Elsevier Inc., 2007).

2. Mostany, R. & Portera-cailliau, C. A Craniotomy Surgery Procedure for Chronic Brain Imaging. J. Vis. Exp. e680 (2008). doi:10.3791/680

3. Siegle, J. H. et al. Open Ephys: An open-source, plugin-based platform for multichannel electrophysiology. J. Neural Eng. 14, 1–13 (2017).

4. Mogensen, F., Müller, D. & Valentin, N. Glycopyrrolate during ketamine/diazepam anaesthesia. A double blind comparison with atropine. Acta Anaesthesiol. Scand. 30, 332–336 (1986).

5. Arena, A. et al. Linear transformation of the encoding mechanism for light intensity underlies the paradoxical enhancement of cortical visual responses by sevoflurane. J. Physiol. 595, 321–339 (2017).

6. Idvall, J., Aronsen, K. F. & Stenberg, P. Tissue Perfusion and Distribution of Cardiac Output During Ketamine Anesthesia in Normovolemic Rats. Acta Anaesthesiol. Scand. 24, 257–263 (1980).

7. Jacobs, G. H., Fenwick, J. A. & Williams, G. A. Cone based vision of rats for ultraviolet and visible lights. J. Exp. Biol. 204, 2439–2446 (2001).

8. De Farias Rocha, F. A. et al. Spectral sensitivity measured with electroretinogram using a constant response method. PLoS One 11, 1–12 (2016).

9. Colombo, M. A. et al. The spectral exponent of the resting EEG indexes the presence of consciousness during unresponsiveness induced by propofol, xenon, and ketamine. Neuroimage 189, 631–644 (2019).

10. Steriade, M., Contreras, D., Amzica, F. & Timofeev, I. Synchronization of fast (30-40 Hz) spontaneous oscillations in intrathalamic and thalamocortical networks. J. Neurosci. 16, 2788–2808 (1996).

11. Mukovski, M., Chauvette, S., Timofeev, I. & Volgushev, M. Detection of active and silent states in neocortical neurons from the field potential signal during slow-wave sleep. Cereb. Cortex 17, 400–414 (2007).

13. Rosanova, M. et al. Sleep-like cortical OFF-periods disrupt causality and complexity in the brain of unresponsive wakefulness syndrome patients. Nat. Commun. 9, 1–10 (2018).

14. Cohen, M. X. Analyzing neural time series data: theory and practice. (MIT Press, 2014).

15. Comolatti, R. et al. A fast and general method to empirically estimate the complexity of brain responses to transcranial and intracranial stimulations. Brain Stimul. 12, 1280–1289 (2019).

16. Marwan, N., Carmen Romano, M., Thiel, M. & Kurths, J. Recurrence plots for the analysis of complex systems. Phys. Rep. 438, 237–329 (2007).

17. Lopes, G. et al. Bonsai: an event-based framework for processing and controlling data streams. Front. Neuroinform. 9, 1–14 (2015).

