## Supplementary Fig. for "General anaesthesia disrupts complex cortical dynamics in response to intracranial electrical stimulation in rats"

### SUPPLEMENTARY MATERIAL

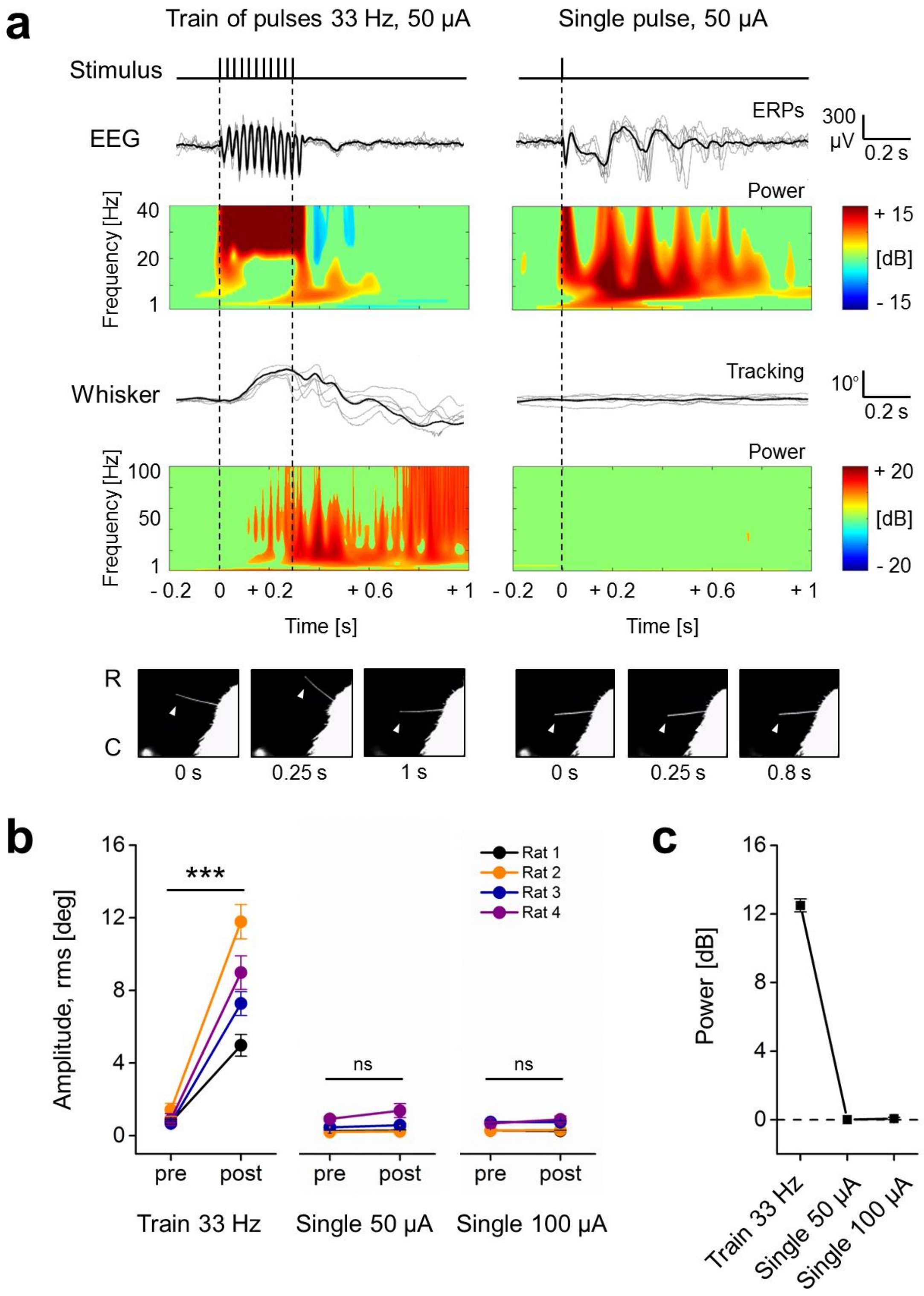

**Supplementary Fig. 1 Single pulse electrical stimulation of M2 did not trigger whisker movements. a**, *up*, example of 5 consecutive ERPs (grey traces) with ensemble average (black) during wakefulness, in response to 2 different electrical stimulations of right M2 (*left*, train of pulses of 1 ms, 50 µA, rate 33 Hz, train duration 0.3 s; *right*, single pulse, 1 ms, 50 µA; 21 stimulations delivered at 0.2 Hz) with relative spectrogram below. *Middle*, tracking of the angular movement of left C1 whisker from the same rat, in response to stimulations (5 consecutive single motor responses in grey and ensemble average in black) with relative spectrogram. *Bottom*, view from above of the rat's snout and C1 whisker (R-C: rostral-caudal) at 3 time points. White arrowheads indicate whisker positions, while the white dot in the bottom-left side represents the onset of the electrical pulse. Only the train stimulation is able to trigger a detectable motor response of the left C1 vibrissae, spanning in a broad frequency range. **b**, The rms amplitude of the angular motor response to the train stimulation and to the single pulse stimulation at 50 and 100 µA is quantified at the level of each animal (n = 4 rats). The rms amplitudes of 0.25 s after the stimulus offset (post) were compared with those obtained from 0.25 s before the onset of stimulation (pre; baseline, from - 0.5 to - 0.25 s) for all the 21 evoked responses (Wilcoxon S-R test; Train stimulation, for all Rats: $P = 5.957*10^{-5}$; Single pulse 50 µA, for Rat 1: $P$ = 0.715, for Rat 2: $P$ = 0.114, for Rat 3: $P$ = 0.170, for Rat 4: $P$ = 0.068; Single pulse 100 µA, for Rat 1: $P$ = 0.931, for Rat 2 and Rat 3: $P$ = 0.664, for Rat 4: $P$ = 0.339). Averaged rms values across rats have been also calculated. Train stimulation: pre stimulus 0.94 ± 0.17°, post stimulus 8.25 ± 1.43°; Single pulse 50 µA: pre stimulus 0.46 ± 0.16°, post stimulus 0.62 ± 0.26°; Single pulse 100 µA:, pre stimulus 0.50 ± 0.12°, post stimulus 0.56 ± 0.16°). **c**, In order to be able to detect also putative small and not phase-locked oscillations of left C1 vibrissae induced by electrical stimulation, a wavelet convolution on whisker tracking was performed. Only the increments or decrements in power for each frequency from 5 to 100 Hz that were statistically different from baseline ($P < 0.05$) have been considered and averaged in a time window of 0.25 s after the offset of stimulation across rats (n = 4). The train of pulses induced a clear increase of power in the broad frequency range (12.49 ± 0.38 dB), but the single pulse stimulations did not trigger any clear power increase as it was close to baseline values (0 ± 0 dB and 0.07 ± 0.07 dB in response to single pules of 50 and 100 µA respectively).

In order to control for the presence of possible body movements induced by the electrical stimulation of M2, a previous set of pilot experiments were also carried out (data not shown). The secondary motor cortex of 2 rats was stimulated by trains of monophasic electrical pulses (50 - 100 µA, 1 ms, delivered at 50 Hz) during the exposure to low dosage of ketamine (1.75 mg/kg/min i.v., rats were responsive to pain stimulations), without body and head restriction. In these conditions only coordinated movements of whiskers (triggered by 86 % of stimulations) and of the head (triggered by 38 % of stimulations) were identified. No other motor responses from other parts of the body could be detected. The M2 cortex of 2 rats was then exposed to monophasic electrical pulses (50 µA, 1 ms) organized in both single pulse stimulations and train of pulses (25 Hz) during wakefulness, in condition of body restriction with free head. While the 100 % of train stimulations triggered a clear coordinated motor response of both head and whiskers (bilaterally), no whisker or head movement could be associated to the single pulse electrical stimulation by visual inspection. We deduced that only train simulations were able to trigger motor responses. The more accurate experiment of video tracking was designed for detecting putative small and fast whisker movements induced by the single pulse electrical stimulation used for replicating PCI relate experiments during wakefulness (here reported).

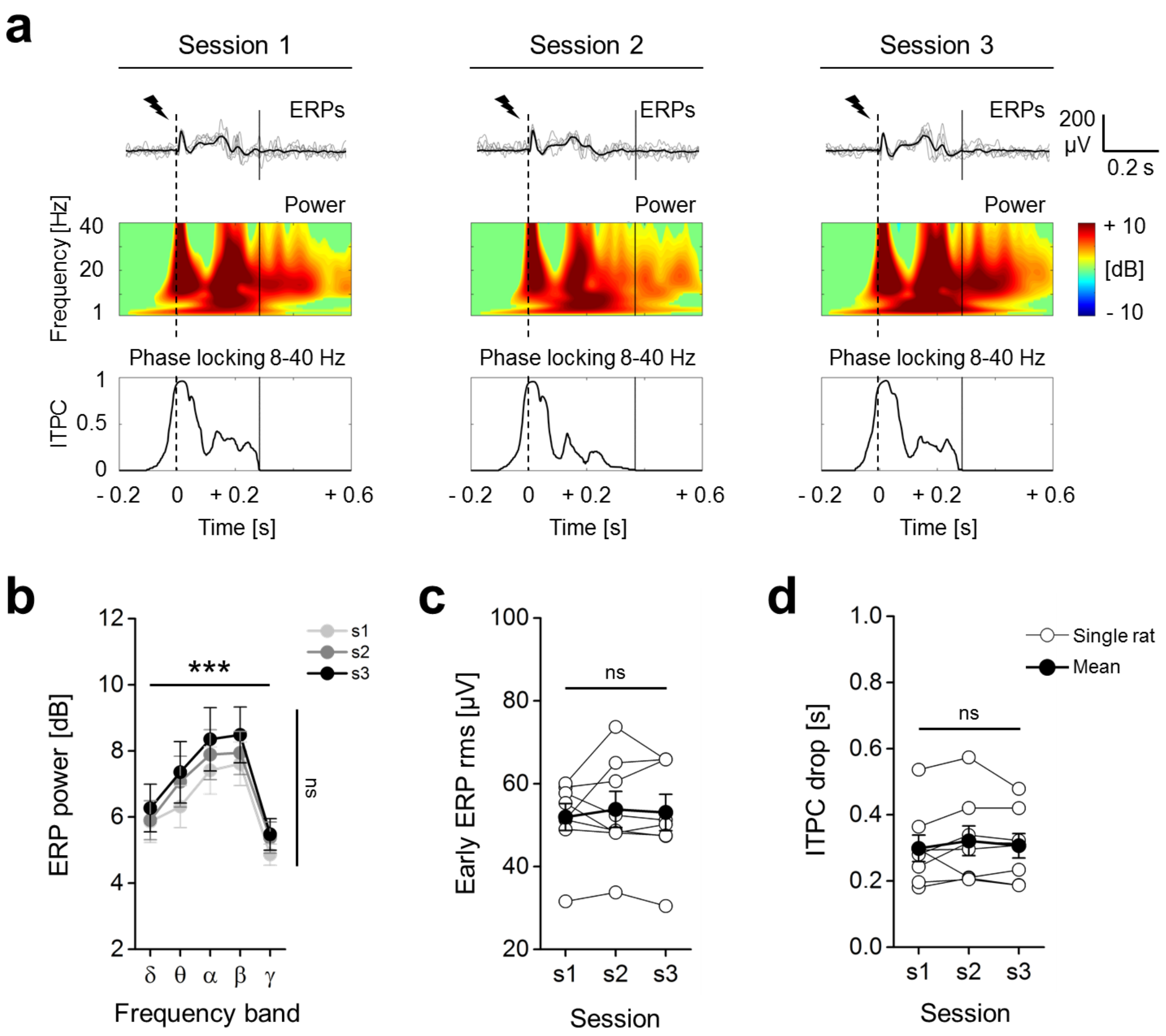
a
Session 1
Session 2
Session 3
ERPs
200 μV
0.2 s
Power
Frequency [Hz]
40
20
1
+ 10
[dB]
- 10
Phase locking 8-40 Hz
ITPC
1
0.5
0
- 0.2
0
+ 0.2
+ 0.6
Time [s]
b
ERP power [dB]
***
ns
s1
s2
s3
δ θ α β γ
Frequency band
c
Early ERP rms [μV]
ns
Session
d
ITPC drop [s]
ns
Single rat
Mean
Session

**Supplementary Fig. 2 Single pulse electrical stimulation of M2 triggered reproducible ERPs that were reliable across different recording sessions. a,** example of 5 consecutive ERPs (grey traces) with ensemble average (black) during wakefulness, in response to single pulse electrical stimulations of M2 (dashed line, 50 µA, 1 ms) from the same rat and channel, in different recording sessions, performed in different days. *Below*, the relative spectrograms are shown, with the inter-trial phase clustering (ITPC) averaged across frequencies (range 8-40 Hz). ITPC quantify the degree of reproducibility of the ERPs (phase-locking across trials). The drop time of ITPC is highlighted by a black line and indicates the duration of the phase-locked response induced by the stimulus (ITPC drop). **b-d**, quantification of the reliability of ERPs in response to single pulse stimulations from n = 8 rats, across 3 recording sessions (s1, s2, s3) performed in different days (∼4 days between each session). **b**, the relative mean spectral power of the ERPs (up to 0.5 s) differed across frequency bands (Friedman test, $P = 1.189*10^{-5}$), with a peak in alpha (8-14 Hz) and beta (15-25 Hz) ranges, but no difference across sessions was identified (Friedman test, *P* = 0.088). **c**, The cortical excitation in response to stimulation was measured as the root mean squared (rms) amplitude of the first deflection of the mean ERPs (Early ERP rms, up to 0.05 s from stimulus onset) and no variation across different days was detected (Friedman test, *P* = 0.687). **d**, Likewise, no significant variation across sessions was detected in the time of ITPC drop (Friedman test, *P* = 0.072).

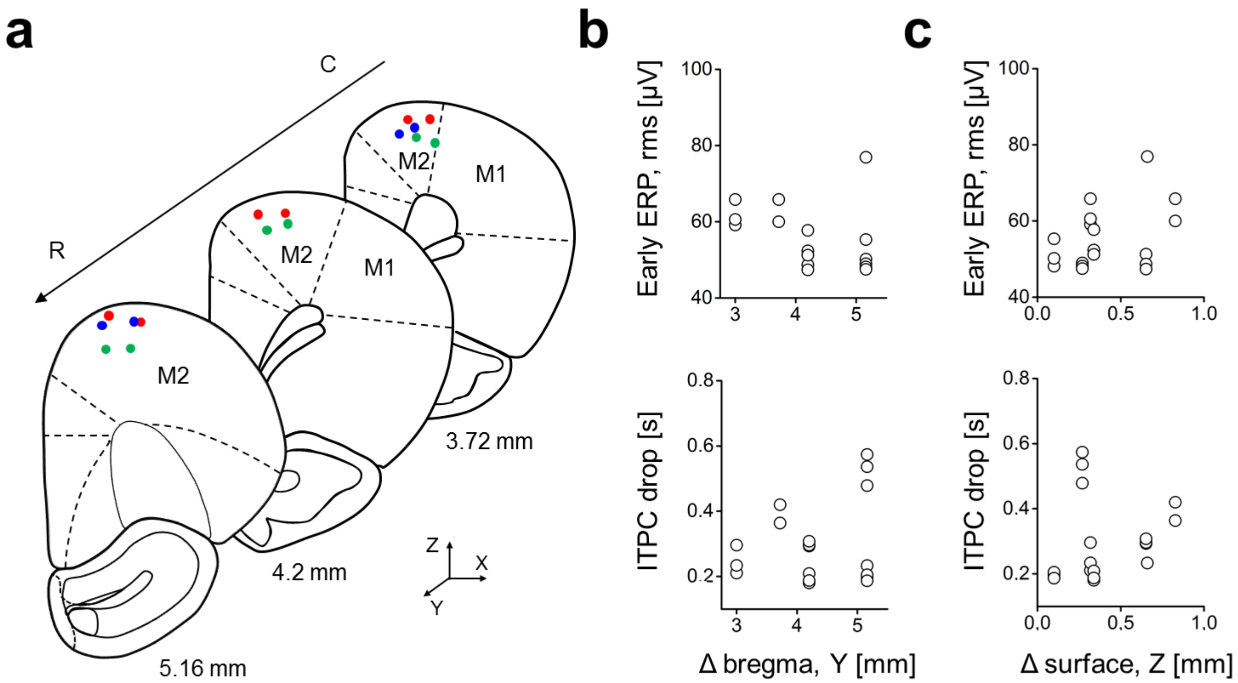
a
C
R
M2
M1
M2
M1
M2
3.72 mm
4.2 mm
5.16 mm
Z
X
Y
b
Early ERP, rms [µV]
ITPC drop [s]
Δ bregma, Y [mm]
c
Early ERP, rms [µV]
ITPC drop [s]
Δ surface, Z [mm]

**Supplementary Fig. 3 Amplitude of ERPs and duration of phase-locked response are not related with the stimulus location in M2. a,** Representation of coronal sections of rat brain with actual position of chronically implanted bipolar electrodes in right secondary motor cortex from 8 rats (dots represent the 2 poles of the bipolar electrode and each rat is colour coded in each slice). Coordinates have been measured from Nissl stained coronal sections of rat brains. Among the tested animals, stimulating electrodes covered a cortical area from ~5 to ~3 mm with respect to bregma in rostro-caudal direction (R-C, Y axes, mean position 4.38 ± 0.25 mm) and a depth range from ~0.1 to ~0.8 mm calculated from cortical surface (Z axes, mean position 0.47 ± 0.09 mm, averaged values for each electrode between the 2 poles), mainly corresponding to layer II/III. On average the 2 electrode poles were separated by 0.46 ± 0.03 mm in the medio-lateral direction (X axes). **b**, **c**, mean rms amplitude of the first deflections of the ERPs (up to 0.05 s from stimulus onset, *up*) and mean duration of phase-locking among subsequent ERPs (ITPC drop, 8-40 Hz, *bottom*) are plotted for all rats (n = 7) and recording sessions (1 to 3 recordings for each rat) during wakefulness against the position of the stimulating electrode along the Y axes (**b**) and along the Z axes (**c**). Possible correlations with the position of stimulating electrode have been tested and no significant relation was detected between the Early ERP rms amplitude and the position of electrodes along the Y axes (**b**, *up;* linear fit, $P = 0.121$, $R^2 = 0.144$) or along the Z axes (**c**, *up;* linear fit, $P = 0.122$, $R^2 = 0{,}143$). No significant relation was identified between the ITPC drop and the position of electrodes along the Y axes (**b**, *bottom;* linear fit, $P = 0.35$, $R^2 = 0{,}055$) or along the Z axes (**c**, *middle;* linear fit, $P = 0.122$, $R^2 = 0{,}143$).

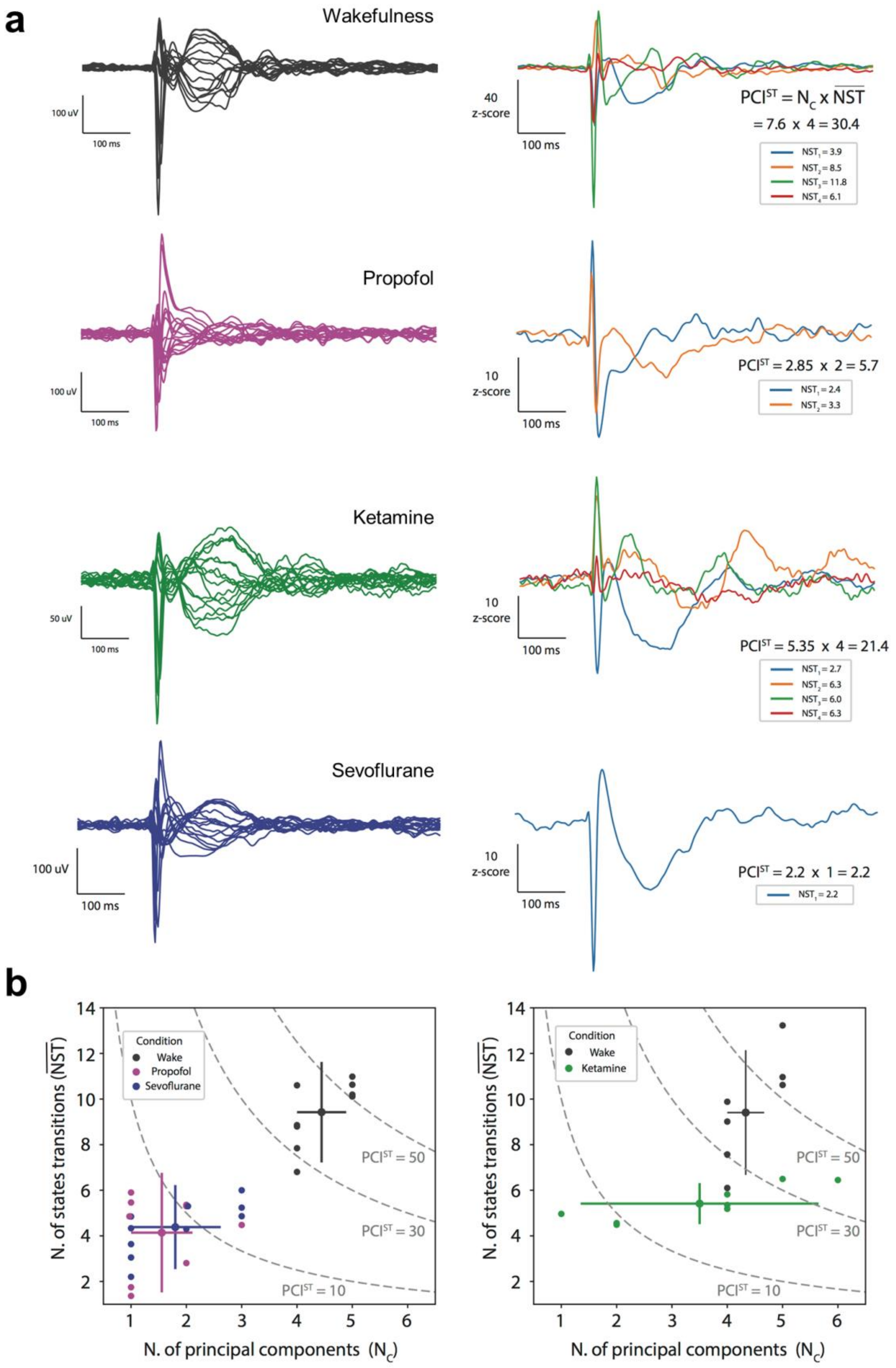
a
Wakefulness
100 uV
100 ms
40
z-score
100 ms
PCI^ST = N_C x NST
= 7.6 x 4 = 30.4
NST1 = 3.9
NST2 = 8.5
NST3 = 11.8
NST4 = 6.1
Propofol
100 uV
100 ms
10
z-score
100 ms
PCI^ST = 2.85 x 2 = 5.7
NST1 = 2.4
NST2 = 3.3
Ketamine
50 uV
100 ms
10
z-score
100 ms
PCI^ST = 5.35 x 4 = 21.4
NST1 = 2.7
NST2 = 6.3
NST3 = 6.0
NST4 = 6.3
Sevoflurane
100 uV
100 ms
10
z-score
100 ms
PCI^ST = 2.2 x 1 = 2.2
NST1 = 2.2
b
Condition
Wake
Propofol
Sevoflurane
PCI^ST = 50
PCI^ST = 30
PCI^ST = 10
N. of states transitions (NST)
N. of principal components (N_C)
Condition
Wake
Ketamine
PCI^ST = 50
PCI^ST = 30
PCI^ST = 10
N. of states transitions (NST)
N. of principal components (N_C)

**Supplementary Fig. 4 PCI$^{ST}$ can be decomposed in average number of state transitions (NST) and number of principal components ($N_C$).** **a**, Event related potentials (ERPs) and corresponding PCI$^{ST}$ decomposition into principal components and state transitions (NST) during wakefulness and during propofol, ketamine and sevoflurane anaesthesia (top to bottom) from one animal. *Left*, butterfly plots with evoked responses to electrical stimulation of right M2. *Right*, PCI$^{ST}$ can be decomposed as the product between the number of principal components ($N_C$), an estimate of the spatial diversity of the signal, and the average number of state transitions (NST), corresponding to the temporal differentiation of the signal, i.e. PCI$^{ST}$ = average NST x NC. The panel shows each principal component of the ERPs that together account for 99% of the variance in the response (time range: 0.08-0.6 s), with the corresponding values of state transitions in the legend box; above it, the PCI$^{ST}$ value computed as the product of average number state transitions (NST) and number of principal components ($N_C$). **b**, Scatter plots with average number state transitions (NST) and number of principal components for all animals with corresponding group average, and 5 and 95 percentiles ($N_C$). *Left*, rats during wake (n = 9), propofol (n = 9) and sevoflurane (n = 10); right, rats during wake (n = 8) and ketamine (n = 8). Also shown for reference are the contour lines (dotted gray lines) with different PCI$^{ST}$ values.

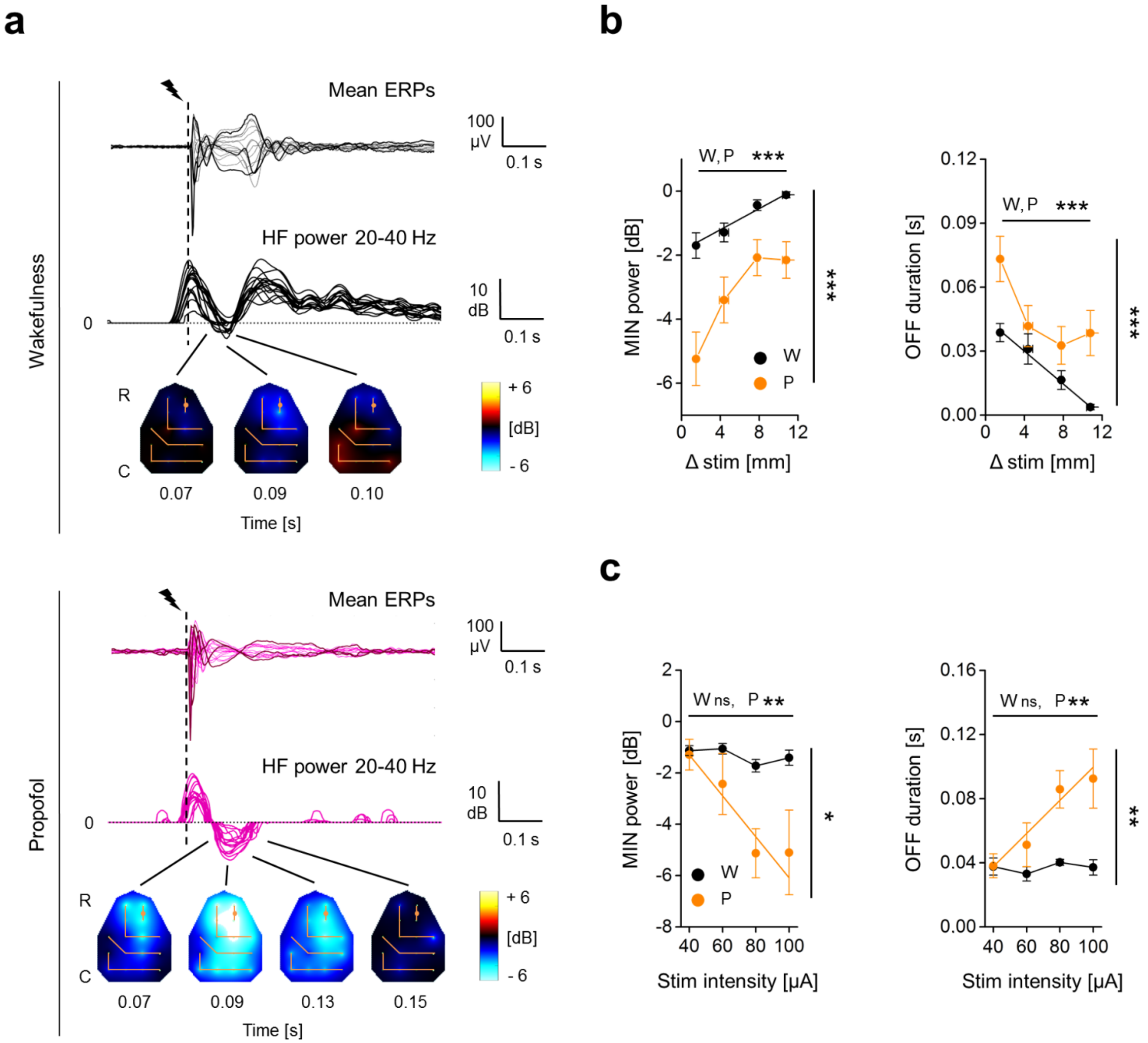
a
b
c
Wakefulness
Propofol
Mean ERPs
HF power 20-40 Hz
100 µV
10 dB
0.1 s
0
R
C
+ 6
[dB]
- 6
0.07
0.09
0.10
0.13
0.15
Time [s]
MIN power [dB]
OFF duration [s]
Δ stim [mm]
Stim intensity [µA]
W, P ***
W ns, P **
W
P
***
*
**

**Supplementary Fig. 5 The suppression of HF power detected during wakefulness was weaker and briefer than the one with propofol and was not affected by increasing stimulus intensity. a,** Epidural EEG activity in response to single pulse electrical stimulation (1 ms, 50 µA; dashed line) of the right secondary motor cortex, from the same rat during wakefulness (*up*) and during propofol anaesthesia (*bottom*). The electrophysiological traces in the butterfly plot represent the superimposition of ensemble averages of ERPs (n = 90 trials) from all recording electrodes (mean ERPs from 3 channels are in bold for highlighting difference in complexity, same channels between wakefulness and propofol conditions). *Below,* the temporal dynamic of HF power (averaged in the range 20-40 Hz) from all the channels is also reported (dotted horizontal line indicates 0 dB). The colour maps show the color-coded interpolation of HF power during the OFF period (window of time when HF power is < 0 dB, suppression) over the scalp in a rostro-caudal orientation (R-C). Orange lines represent the isodistance lines that connect channels with similar spatial distance from the stimulating electrode (orange dot, M2 right). To assess the relation between the OFF period and the spatial distance from the stimulus, we averaged values from channels that were clustered in 4 isodistance lines, at 1.5 ± 0 mm, 4.38 ± 0.47 mm, 7.8 ± 0.17 mm and 10.8 ± 0.40 mm far from the site of stimulation. **b**, Quantification of the negative peak of HF power during the OFF period (deepest suppression, MIN power, *left*) and duration of the suppression of HF power (OFF duration, *right*), averaged within isodistance lines and across rats (n = 9 animals) during wakefulness (W) and propofol anaesthesia (P). During wakefulness, MIN power linearly approaches to 0 dB from more negative values while increasing the distance from the site of stimulation (Friedman test, $P = 6.022*10^{-5}$; linear fit, $P = 0.021$, $R^2 = 0.959$). Also during propofol anaesthesia a relation between MIN power and distance from stimulation could be detected (Friedman test, $P = 8.694*10^{-5}$), but this was not linear (linear fit, $P = 0.103$, $R^2 = 0.805$). Overall, MIN power was more close to 0 dB during wakefulness then during propofol anaesthesia (Friedman test, $P = 4.817*10^{-6}$). Likewise, OFF duration linearly decreased approaching 0 s by increasing the distance from the stimulus site during wakefulness (Friedman test, $P = 2.458*10^{-5}$; linear fit, $P = 0.001$, $R^2 = 0.997$). A relation between OFF duration and distance from stimulus could be also detected during propofol anaesthesia, but this was not linear (Friedman test, $P = 2.734*10^{-4}$; linear fit, $P = 0.213$, $R^2 = 0.618$). Overall, OFF duration was shorter during wakefulness then during propofol anaesthesia (Friedman test, $P = 2.205*10^{-4}$). **c**, In addition, both MIN power (*left*) and OFF duration (*right*, averaged across all channels) were found to change differently in relation to the intensity of stimulation between wakefulness and propofol anaesthesia in n = 5 rats. No significant trends were found during wakefulness (MIN power, Friedman test, $P = 0.145$; OFF duration, Friedman test, $P = 0.472$).

Otherwise, during propofol anaesthesia, MIN power significantly decreased by increasing intensity of stimulation with a trend that could be linearly fitted (Friedman test, $P$ = 0.007; linear fit, $P$ = 0.035, $R^2$ = 0.932) and OFF duration linearly increased by increasing stimulus intensity (same data from Fig. 3; Friedman test, $P$ = 0.005; linear fit, $P$ = 0.024, $R^2$ = 0.953). Overall, also in relation to stimulus intensity, MIN power was lower and OFF duration was higher during propofol anaesthesia than during wakefulness (MIN power, Friedman test, $P$ = 0.037; OFF duration, Friedman test, $P$ = 0.003).

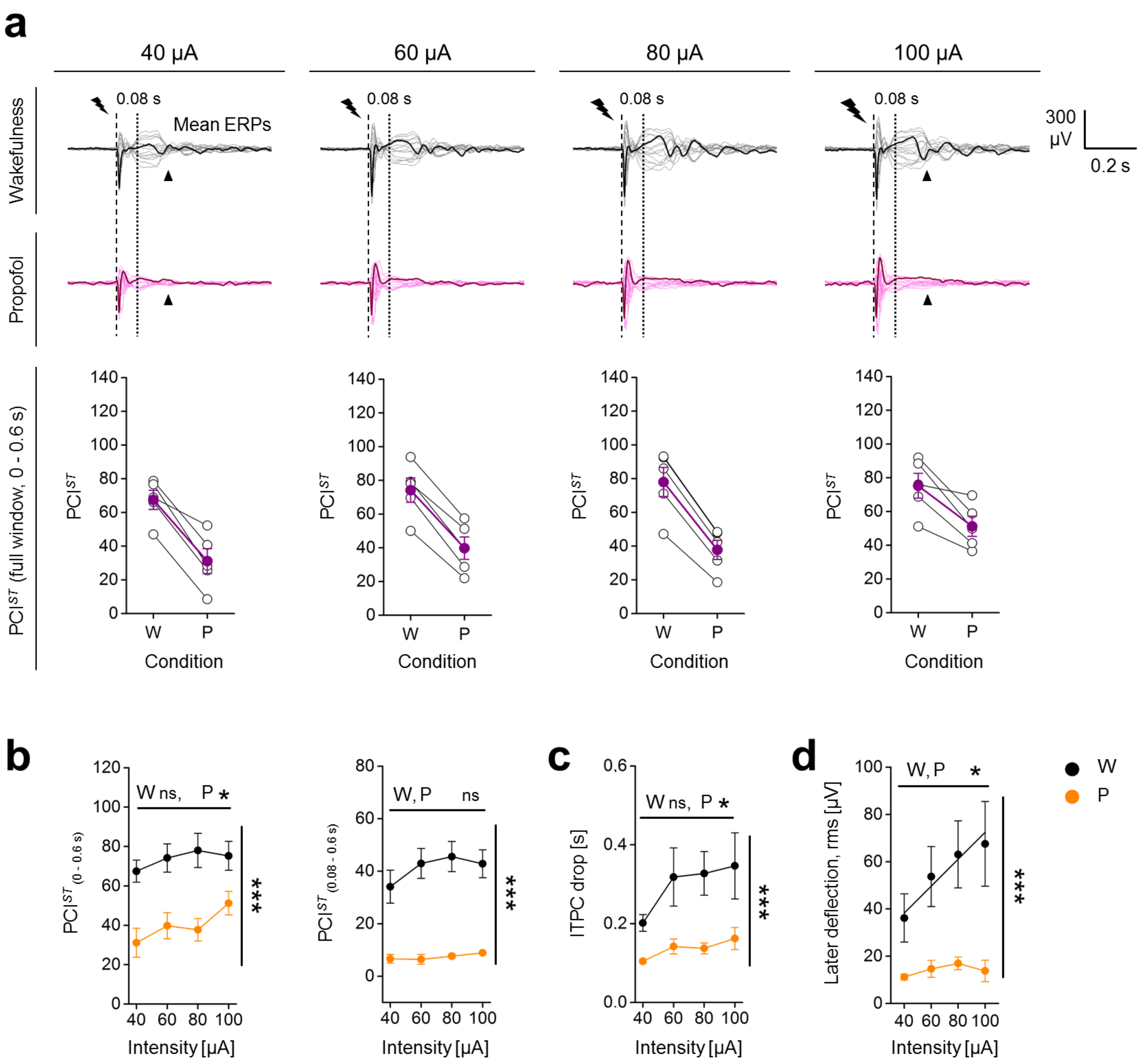

a
40 μA
60 μA
80 μA
100 μA
Wakefulness
Propofol
0.08 s
Mean ERPs
300 μV
0.2 s
PCI$^{ST}$ (full window, 0 - 0.6 s)
PCI$^{ST}$
W
P
Condition
b
PCI$^{ST}$ (0 - 0.6 s)
PCI$^{ST}$ (0.08 - 0.6 s)
W ns, P *
W, P ns
***
Intensity [μA]
c
ITPC drop [s]
W ns, P *
d
Later deflection, rms [μV]
W, P *
W
P

**Supplementary Fig. 6 Perturbational complexity in relation to incremental intensity of stimulation. a,** epidural EEG activity in response to single pulse electrical stimulations of increasing intensities (1 ms, 40, 60, 80 and 100 µA; dashed line) of the right secondary motor cortex from the same rat during wakefulness (*up*) and during the exposure to propofol anaesthesia (*middle*). The electrophysiological traces represent the superimposition of ensemble averages of evoked related potentials from all the 16 recording electrodes distributed all over the skull. One averaged ERP from the same channel across condition has been highlighted in bold for better illustrating the difference in cortical complexity. The vertical dotted lines indicate 0.08 s, which approximates the rising of the OFF period. $PCI^{ST}$ scores obtained from the whole window of the evoked response (0-0.6 s) are shown for all animals (white dots) and averaged across rats (n = 5 rats; purple dots) for all intensities of stimulation and condition (*below*). $PCI^{ST}$ always decreased in all animals, from wakefulness (W) to propofol anaesthesia (P), regardless the intensity of stimulation. **b**, We otherwise observed a significant variation of $PCI^{ST}$ (in full time window 0-0.6 s; *left*) during propofol anaesthesia in relation to stimulus intensity (Friedman test, $P$ = 0.029), which was not detectable during wakefulness (Friedman test, $P$ = 0.062). Overall $PCI^{ST}$ during wakefulness was higher than during propofol anaesthesia (Friedman test, $P = 1.55*10^{-5}$). No relation with stimulus intensity was detected by calculating $PCI^{ST}$ from 0.08 to 0.6 s and therefore by excluding the first response to the electrical stimulation, before the rising of the OFF period (*right*). No significant variation of $PCI^{ST}$ has been observed during propofol anaesthesia (Friedman test, $P$ = 0.782), but a dim variation seemed to be present during wakefulness even if did not reached statistical significance (Friedman test, $P$ = 0.0503). Overall perturbational complexity during wakefulness was higher than during propofol anaesthesia (Friedman test, $P = 1.767*10^{-7}$). **c**, Quantification of ITPC drop time (8-40 Hz; *right*) as a function of increasing intensity of stimulation during both wakefulness and propofol anaesthesia. Values are averaged across channels and animals. No statistically significant variation was detected in wakefulness condition (Friedman test, $P$ = 0.077). Otherwise, during propofol anaesthesia, we identified a significant variation of ITPC drop time (Friedman test, $P$ = 0.02) as a function of stimulus intensity. This was in line with the correlation observed with the end of the OFF period (see Fig. 3). Overall, ITPC drop time values obtained during wakefulness were higher than during propofol anaesthesia (ITPC drop, Friedman test, $P = 7.082*10^{-6}$). **d**, The quantification of the amplitude of a later deflections of ERPs in relation to stimulus intensity, between wakefulness and anaesthesia is reported. RMS amplitude has been measured around 0.2 s (in time window 0.175-0.225 s, black arrowheads in **a**) of the mean ERP from all channels and then averaged for each animal (n = 5 rats) and condition. As for the early evoked response (see Fig. 3), a linear increase of the amplitude of the later deflection of ERPs has been detected as a function of stimulus intensity in wakefulness (Friedman test, $P$ = 0.013; linear fit, $P$ = 0.032, $R^2$ = 0.936). Also in propofol anaesthesia the amplitude of the later deflection of the ERPs changed as a function of stimulus intensity, but it was not linear (Friedman test, $P$ = 0.04; linear fit, $P$ = 0.14, $R^2$ = 0.739). Overall, differently from the early evoked response (Fig. 3), the amplitude of the later deflection of ERPs has been found to be significantly higher in wakefulness than during propofol anaesthesia (Friedman test, $P = 1.767*10^{-7}$).

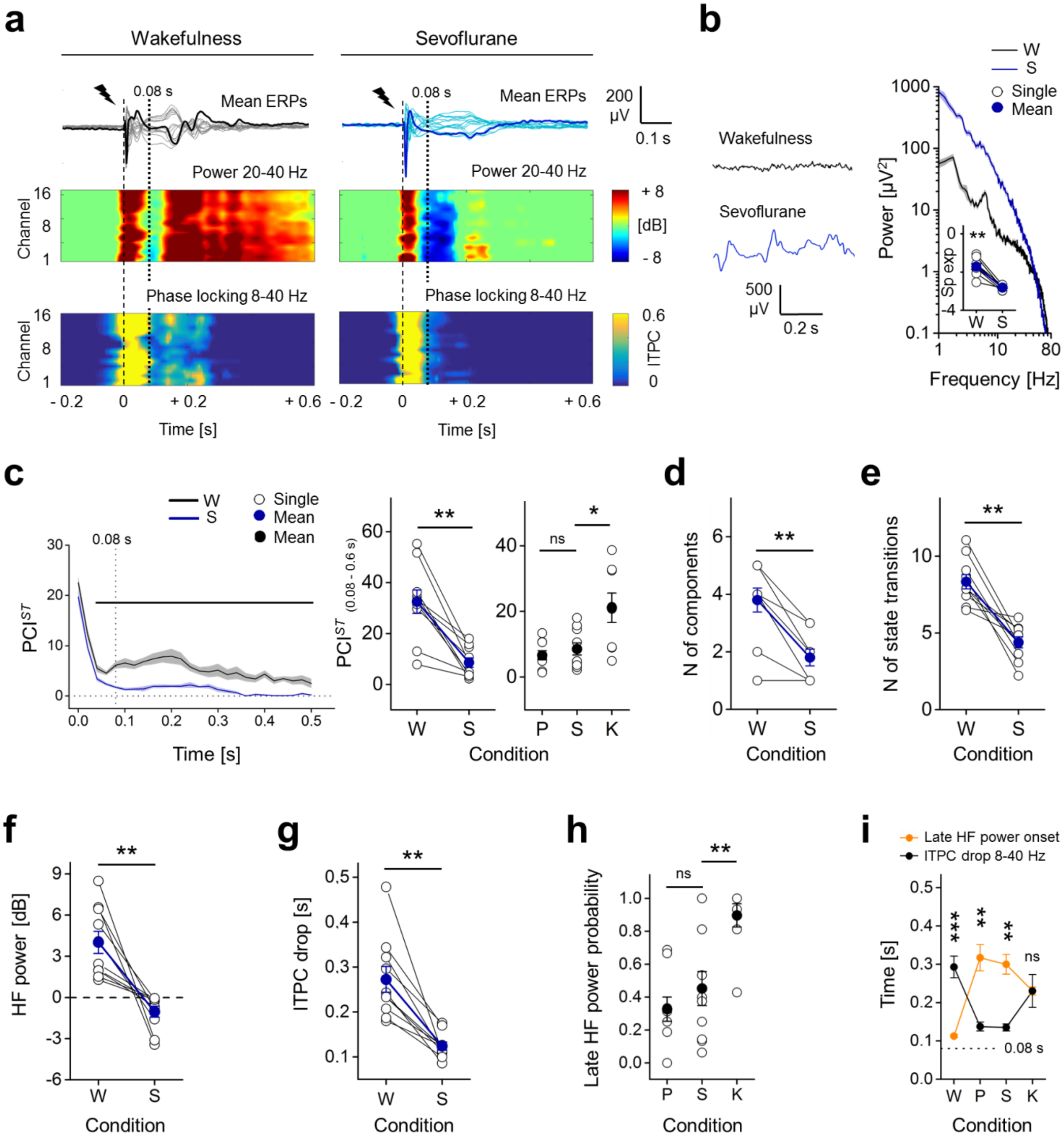

a
Wakefulness
Sevoflurane
0.08 s
Mean ERPs
200 µV
0.1 s
Power 20-40 Hz
Channel
+ 8
[dB]
- 8
Phase locking 8-40 Hz
0.6
ITPC
0
- 0.2
0
+ 0.2
+ 0.6
Time [s]
b
Wakefulness
Sevoflurane
500 µV
0.2 s
W
S
Single
Mean
Power [µV²]
Sp exp
Frequency [Hz]
c
PCI$^{ST}$
PCI$^{ST}$ (0.08 - 0.6 s)
Time [s]
Condition
ns
d
N of components
e
N of state transitions
f
HF power [dB]
g
ITPC drop [s]
h
Late HF power probability
i
Late HF power onset
ITPC drop 8-40 Hz
Time [s]
W P S K

**Supplementary Fig. 7 PCI$^{ST}$, suppression of high frequencies and phase-locking of ERPs in response to electrical stimulation during sevoflurane anaesthesia. a,** Superimposition of mean ERPs from all electrodes in response to single pulse stimulation (1 ms, 50 µA; dashed line) from the same rat during wakefulness (W) and sevoflurane (S; ~2.6 %) anaesthesia (same rat from Fig. 4 to facilitate comparisons). One averaged ERP from the same channel (M2) is in bold to highlight differences in complexity. Spectrograms of HF power and ITPC for all channels are shown below. Vertical dotted line at 0.08 s indicates the average time of OFF period onset across rats. **b**, Spontaneous EEG (*left*) and relative mean periodograms (shades represent SEM; *right*) are reported from one rat during wakefulness and sevoflurane anaesthesia. Spectral exponents from all rats are also reported (*inset*). The spectral exponent dropped from wakefulness to sevoflurane anaesthesia, highlighting a redistribution of power towards slow frequencies (n = 9 rats; wakefulness: -1.71 ± 0.15, sevoflurane: -2.81 ± 0.04; Wilcoxon S-R test, $P$ = 0.004). **c,** *Left*, time courses of mean PCI$^{ST}$ in conditions of wakefulness and sevoflurane anaesthesia (n = 10 rats; shades represent SEM; horizontal lines indicate time periods with statistical differences between conditions, Wilcoxon S-R test $P < 0.05$). *Right*, PCI$^{ST}$ in range 0.08-0.6 s is reported for each rat and condition. PCI$^{ST}_{(0.08\text{-}0.6\ s)}$ significantly dropped from wakefulness to sevoflurane anaesthesia (wakefulness: 32.50 ± 4.60, sevoflurane: 8.57 ± 1.85; Wilcoxon S-R test, $P$ = 0.002). Moreover with sevoflurane, PCI$^{ST}_{(0.08\text{-}0.6\ s)}$ was similar to what obtain with propofol, but significantly lower compared to ketamine condition (Mann-Whitney test; propofol vs sevoflurane, $P$ = 0.775; ketamine vs sevoflurane, $P$ = 0.018). Like for propofol anaesthesia, the decreased PCI$^{ST}_{(0.08\text{-}0.6\ s)}$ compared to wakefulness was explained by both a reduced number of principal components (**d**, wakefulness: 3.80 ± 0.42, sevoflurane: 1.80 ± 0.29; Wilcoxon S-R test, $P$ = 0.004) and reduced averaged number of state transitions (**e**, wakefulness: 8.33 ± 0.47, sevoflurane: 4.38 ± 0.36; Wilcoxon S-R test, $P$ = 0.002) of the EEG response to the stimulation. **f**, During sevoflurane anaesthesia, the electrical stimulation triggered a first response followed by an OFF period in all animal tested starting at 0.075 ± 0.005 s (average across channels and rats), characterized by a profound suppression of HF power (20-40 Hz). By averaging HF power across channels, in time range 0.08-0.18 s, a significant difference from wakefulness was detected (wakefulness: 4.01 ± 0.81 dB, sevoflurane: -1.04 ± 0.40 dB, Wilcoxon S-R test, $P$ = 0.002). **g,** The mean ITPC drop time (8-40 Hz) across channels also significantly decreased from wakefulness to sevoflurane anaesthesia (wakefulness: 0.27 ± 0.03 s, sevoflurane: 0.12 ± 0.01 s; Wilcoxon S-R test, $P$ = 0.002) indicating shorter phase-locked ERPs in the latter condition. **h**, With sevoflurane, the ratio between the amount of channels that presented a later increase in HF power and the total number of electrodes (late HF power probability, 0.45 ± 0.10) was similar to what observed with propofol, and significantly lower than during ketamine anaesthesia (Mann-Whitney test, sevoflurane vs propofol, $P$ = 0.46; sevoflurane vs ketamine, $P$ = 0.006). **i**, Timings of late HF power onset and ITPC drop (8-40 Hz) are shown for all conditions. Like for propofol anaesthesia, with sevoflurane the increment of HF power occurred later then ITPC drop time in 8-40 Hz range (Wilcoxon S-R test, sevoflurane $P$ = 0.002), thus indicating a not phase-locked activity. Propofol and ketamine data are the same from Fig. 4, reported here to allow comparisons.

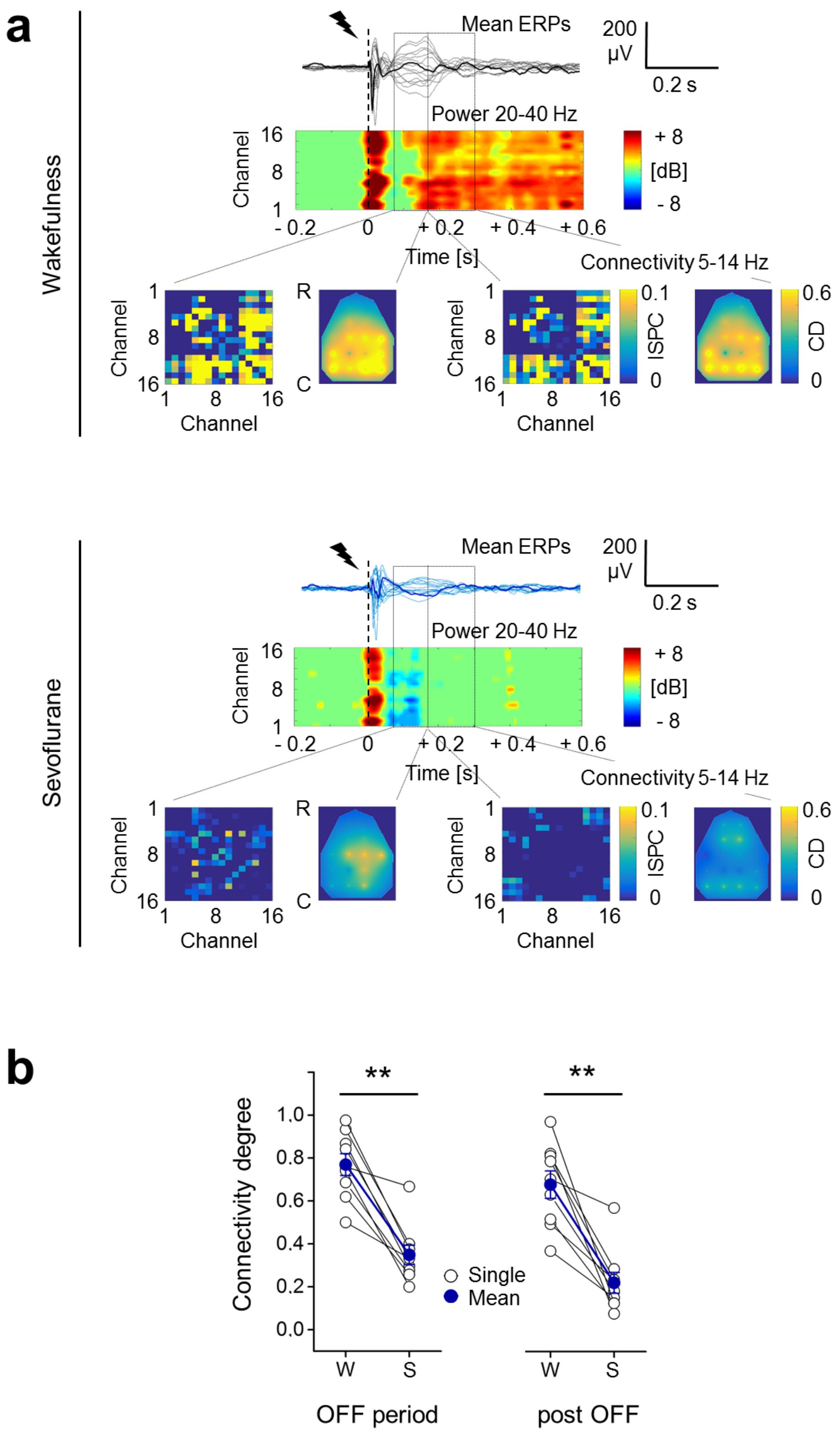
a
Wakefulness
Mean ERPs
200 µV
0.2 s
Power 20-40 Hz
Channel
16
8
1
+ 8
[dB]
- 8
- 0.2
0
+ 0.2
+ 0.4
+ 0.6
Time [s]
Connectivity 5-14 Hz
R
C
ISPC
0.1
0
CD
0.6
Sevoflurane
b
**
Connectivity degree
1.0
0.8
0.6
0.4
0.2
0.0
Single
Mean
W
S
OFF period
post OFF

**Supplementary Fig. 8 Functional connectivity after cortical perturbation during sevoflurane anaesthesia. a,** Example of epidural EEG activity in response to electrical stimulation (1 ms, 50 µA; dashed lines) from the same rat during wakefulness and during sevoflurane anaesthesia (same rat of Fig. 5 to allow comparisons). The electrophysiological traces represent the superimposition of ensemble averages of ERPs from all electrodes. 1 channel is highlighted in bold for better illustrating difference in complexity. Below each butterfly plot, the spectrogram of mean relative HF power (20-40 Hz) is shown for each channel. The bottom part of each inner panel shows the increments in cortical connectivity compared to baseline for each condition in two time windows: OFF period, 0.08-0.18 s, *left* and post OFF, 0.18-0.3 s, *right* (rectangles indicate the two time windows). For each window, the connectivity matrix based on "inter-site phase clustering" (ISPC, 5-14 Hz) is reported on the *left* and the topographical distribution (rostro-caudal orientation, R-C) of the connectivity degree (CD) for each channel is interpolated and shown on the *right*. **b**, Mean CD across channels in the OFF period, *left* and post OFF, *right,* during wakefulness and sevoflurane anaesthesia (n = 9 rats). During wakefulness CD was significantly higher then what observed during sevoflurane anaesthesia in both time windows (OFF period, wakefulness: 0.77 ± 0.05, sevoflurane: 0.35 ± 0.04; post OFF, wakefulness: 0.67 ± 0.06, sevoflurane: 0.23 ± 0.05; Wilcoxon S-R test, for both time windows, $P$ = 0.004).

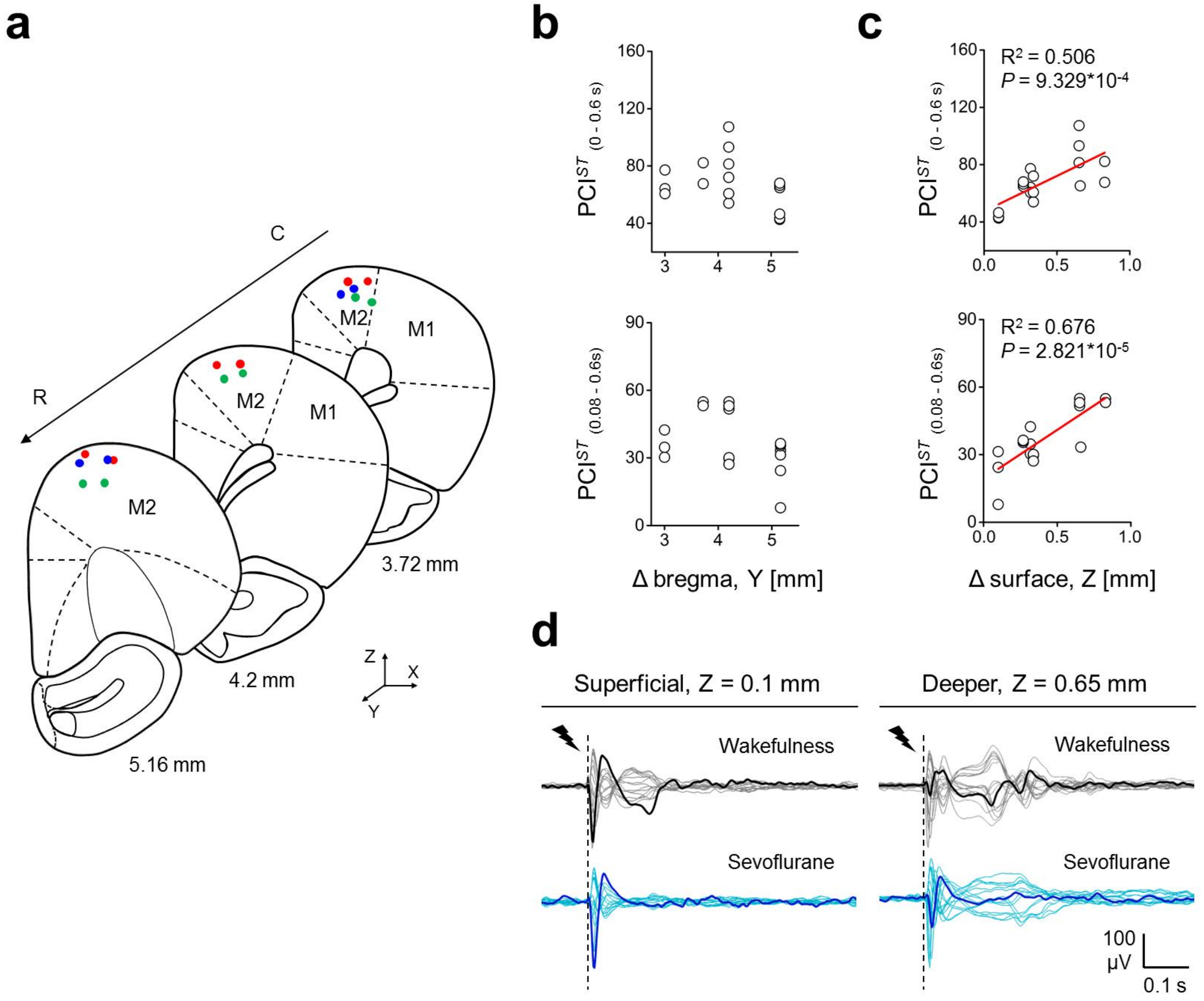
a
C
R
M2
M1
M2
M1
M2
3.72 mm
4.2 mm
5.16 mm
Z
X
Y
b
c
R² = 0.506
P = 9.329*10⁻⁴
R² = 0.676
P = 2.821*10⁻⁵
PCI^ST (0 - 0.6 s)
PCI^ST (0.08 - 0.6s)
Δ bregma, Y [mm]
Δ surface, Z [mm]
d
Superficial, Z = 0.1 mm
Deeper, Z = 0.65 mm
Wakefulness
Sevoflurane
100 μV
0.1 s

**Supplementary Fig. 9 PCI$^{ST}$ correlated with the stimulus location in M2 along the dorso-ventral axis during wakefulness. a**, The same representation of coronal sections of rat brain with the actual position of chronically implanted bipolar electrodes in right M2 has already been shown in Supplementary Fig. 3 and is reported here for clarity. Dots represent the 2 poles of the bipolar electrodes chronically implanted in 8 rats (for each section, bipolar electrodes from different rats are colour coded). Coordinates have been measured from Nissl stained coronal sections of rat brains and the stimulating electrodes were found to cover a cortical area from ~5 to ~3 mm with respect to bregma in rostro-caudal direction (R-C, Y axis, mean position 4.38 ± 0.25 mm) and a depth range from ~0.1 to ~0.8 mm calculated from cortical surface (Z axis, mean position 0.47 ± 0.09 mm, averaged values for each electrode between the 2 poles), mainly corresponding to layer II/III. In one rat the electrode was found to be placed in layer I, while in another animal it was positioned at the edge between layer III and V. **b**, **c**, PCI$^{ST}$ from time range 0-0.6 s (*up*) and from time window 0.08-0.6 s (*bottom*) are plotted for each rat (n = 7) and recording session during wakefulness (1 to 3 recordings for each rat) against the position of the stimulating electrode along the Y axis (**b**) and along the Z axis (**c**). Possible correlations with the position of stimulating electrode have been tested and no significant relation was detected between PCI$^{ST}$ and the position of stimulating electrodes along the Y axes (**b**; *up*, $PCI^{ST}_{(0\text{-}0.6\ s)}$, linear fit, $P = 0.155$, $R^2 = 0.122$; *bottom*, $PCI^{ST}_{(0.08\text{-}0.6\ s)}$, linear fit, $P = 0.129$, $R^2 = 0.138$). Otherwise, a highly significant and strong positive correlation between PCI$^{ST}$ (in both time windows) and the depth of the stimulation site have been found (**c**; *up*, $PCI^{ST}_{(0\text{-}0.6\ s)}$, linear fit, $P = 9.329*10^{-4}$, $R^2 = 0.506$; *bottom*, $PCI^{ST}_{(0.08\text{-}0.6\ s)}$, linear fit, $P = 2.821*10^{-5}$, $R^2 = 0.676$). The coefficient of determination $R^2$ and the $P$ value are reported. **d**, Example of superimposed evoked activity from one rat with stimulating electrode implanted close to cortical surface (*left*) and from another animals with stimulating electrode placed deeper in M2 (*right*), during wakefulness (*up*) and sevoflurane anaesthesia (*bottom*). The electrophysiological traces represent the superimposition of mean ERPs from all recording channels. One mean ERP from the same channel (S1) is in bold in each condition to better highlight differences in complexity. Data are from the same animals reported in Fig. 6.

**Supplementary Video 1** **Train of electrical pulses to M2 triggered whisker movements.** The view from above shows an example of the angular movements of the left C1 whisker from the snout of a representative rat (same rat of Supplementary Video 2) in response to a train of electrical pulses (pulse amplitude 50 µA, pulse duration 1 ms, rate of pulses 33 Hz, train duration 0.3 s) delivered to M2. The lighting of an LED in the upper right part of the panel indicates the onset of the electrical stimulation. It can be noted how in response to the train stimulation a clear and wide angular movement is performed by the left C1 vibrissae.

**Supplementary Video 2** **Single electrical pulse stimulation of M2 did not trigger whisker movements.** The view from above shows an example of the angular movements of the left C1 whisker from the snout of a representative rat (same rat of Supplementary Video 1) in response to single electrical pulse stimulation (pulse amplitude 50 µA, pulse duration 1 ms) delivered to M2. The lighting of a LED in the upper right part of the panel indicates the onset of the electrical stimulation. It can be noted how no clear motor reaction of the left C1 vibrissae can be detected in response to single pulse stimulation.

**Supplementary Video 3** **Spatiotemporal dynamic of EEG response to single pulse electrical stimulation during wakefulness.** Example of epidural EEG activity in response to single pulse electrical stimulation (1 ms, 50 µA; triggered at 0 s) of right M2 from the same rat of Supplementary Video 4, 5 and 6 during wakefulness. On the left, the electrophysiological traces in the butterfly plots represent the superimposition of ensemble averages of ERPs (n = 90 trials) from all 16 recording electrodes (scale of Y axes: µV, scale of X axes: s). On the right, the colour map shows the spatial distribution of the electrodes (circles) over the scalp in a rostro (anterior)-caudal orientation (scale of the Y and X axes: mm ) and the color-coded interpolation of the evoked potentials for single time points (scale bar in µV). The moving red vertical bar in the butterfly plot indicates the time instant represented in the colour map. During wakefulness the evoked response was composed of long-lasting complex waveforms with multiple changes in polarity along time and across cerebral areas.

**Supplementary Video 4 Spatiotemporal dynamic of EEG response to single pulse electrical stimulation during propofol anaesthesia.** Example of epidural EEG activity in response to single pulse electrical stimulation (1 ms, 50 µA; triggered at 0 s) of right M2 from the same rat of Supplementary Video 3, 5 and 6 during propofol anaesthesia. On the left, the electrophysiological traces in the butterfly plots represent the superimposition of ensemble averages of ERPs (n = 90 trials) from all 16 recording electrodes (scale of Y axes: µV, scale of X axes: s). On the right, the colour map shows the spatial distribution of the electrodes (circles) over the scalp in a rostro (anterior)-caudal orientation (scale of the Y and X axes: mm ) and the color-coded interpolation of the evoked potentials for single time points (scale bar in µV). The moving red vertical bar in the butterfly plot indicates the time instant represented in the colour map. During propofol anaesthesia the evoked response was composed of short-lasting waveforms with few polarity changes along time and across cerebral areas. In contrast to wakefulness, after a first response, the peak of voltage was found to be relatively stationary close to the stimulation site (up-right part of the colour map).

**Supplementary Video 5 Spatiotemporal dynamic of EEG response to single pulse electrical stimulation during ketamine anaesthesia.** Example of epidural EEG activity in response to single pulse electrical stimulation (1 ms, 50 µA; triggered at 0 s) of right M2 from the same rat of Supplementary Video 3, 4 and 6 during ketamine anaesthesia. On the left, the electrophysiological traces in the butterfly plots represent the superimposition of ensemble averages of ERPs (n = 90 trials) from all 16 recording electrodes (scale of Y axes: µV, scale of X axes: s). On the right, the colour map shows the spatial distribution of the electrodes (circles) over the scalp in a rostro (anterior)-caudal orientation (scale of the Y and X axes: mm ) and the color-coded interpolation of the evoked potentials for single time points (scale bar in µV). The moving red vertical bar in the butterfly plot indicates the time instant represented in the colour map. During ketamine anaesthesia the evoked response was closer to what seen in wakefulness. It was composed of long-lasting waveforms with some polarity changes along time and across cerebral areas.

**Supplementary Video 6 Spatiotemporal dynamic of EEG response to single pulse electrical stimulation during sevoflurane anaesthesia.** Example of epidural EEG activity in response to single pulse electrical stimulation (1 ms, 50 µA; triggered at 0 s) of right M2 from the same rat of Supplementary Video 3, 4 and 5 during sevoflurane anaesthesia. On the left, the electrophysiological traces in the butterfly plots represent the superimposition of ensemble averages of ERPs (n = 90 trials) from all 16 recording electrodes (scale of Y axes: µV, scale of X axes: s). On the right, the colour map shows the spatial distribution of the electrodes (circles) over the scalp in a rostro (anterior)-caudal orientation (scale of the Y and X axes: mm ) and the color-coded interpolation of the evoked potentials for single time points (scale bar in µV). The moving red vertical bar in the butterfly plot indicates the time instant represented in the colour map. During sevoflurane anaesthesia the evoked response was similar to what obtain with propofol. It was composed of short-lasting waveforms with few polarity changes along time and across cerebral areas. In contrast to wakefulness, after a first response, the peak of voltage was found to be relatively stationary close to the stimulation site (up-right part of the colour map).
